# Language network functional connectivity in infancy predicts developmental language trajectories

**DOI:** 10.1101/2025.09.17.676930

**Authors:** Lauren Wagner, Joshua Ceballos, Emily Chiem, Mirella Dapretto

## Abstract

Although developmental language delays affect approximately 10% of children in the general population, the neurodevelopmental mechanisms that support normative language acquisition, and atypicalities that may predict later language delay, across the first year of life are poorly understood. Here, resting-state fMRI data from the Baby Connectome Project was used to evaluate age-related changes in language network functional connectivity and alterations associated with suboptimal language development. Additionally, a data-driven machine learning algorithm was used to partition our sample into three groups who showed Delayed, Typical, and Advanced trajectories of language development. These groups reliably differed on several assessments of language ability during infancy and toddlerhood. Using *a priori* brain regions involved in adult language processing, a seed-based functional connectivity analysis showed broad age-related increases in functional synchrony and specialization throughout the infant language network. Additionally, the Delayed group showed several atypical patterns of functional connectivity with language regions. Importantly, the magnitude of connectivity differences consistently predicted later language scores at two-year outcome across several different language assessments. These findings add to our understanding of normative neurodevelopmental patterns underlying language acquisition, and identify several potential biomarkers associated with language delay that could serve as future targets to inform diagnoses and clinical interventions.

**Highlights:**

- The language network undergoes significant maturational changes during infancy
- With age, functionally similar nodes integrate, while dissimilar nodes segregate
- Language-delayed infants show atypical connectivity in the language network
- These early atypicalities predict later language development at two-year outcome
- We identify several neural signatures associated with early language delay

## 1. Introduction

Developmental language impairments are common in the general population, affecting an estimated 1 in 10 children (Norbury et al., 2016). These challenges are associated with issues with emotional and behavioral regulation (Yew & O’Kearney, 2013), as well as lower educational attainment and social competence (Tomblin, 2008), and worse employment outcomes (Conti & Durkin, 2012). Moreover, prospective studies in children who later develop a reading disorder such as dyslexia show that early impairments in receptive vocabulary, syntax, and phonemic awareness predate the onset of dyslexia (Scarborough, 1990). However, despite the high rate of occurrence of language and reading disorders and their far-reaching impacts throughout adolescence and young adulthood, very little is currently known about the neurodevelopmental mechanisms that support early language acquisition, both in normative and neurodiverse samples. Early detection and intervention are important for mitigating delayed or impaired language (Almost & Rosenbaum, 1998; Capone Singleton, 2018). Therefore, understanding the neural underpinnings of normative language development may inform earlier detection and timelier interventions that would lead to better prognoses and long-term quality of life.

The first year of life is a critical epoch for language acquisition, as most major developmental milestones supporting the emergence of both speech and comprehension are observed during this period. These behavioral milestones, which include canonical babbling, rapid improvements in joint attention and theory of mind, and the acquisition of a small receptive vocabulary, are relatively consistent across languages, cultures, and even spoken vs. signed modalities (Kern et al., 2009; Mayberry & Squires, B., 2006; Petitto & Marentette, 1991). This behaviorally dynamic period of life is also characterized by dramatic growth in the young brain’s functional architecture. Primary sensory networks are first to resemble adult networks, while higher-order networks develop more gradually (Gao et al., 2015). The more protracted development of higher-order networks suggests that their developmental course could be malleable to intervention. Growing evidence suggests that functional networks involved in language may be disrupted in disorders involving language impairment, such as autism spectrum disorder (ASD; Emerson et al., 2017; Liu et al., 2020; Nair et al., 2021), which may lead to poorer language outcomes (Dinstein et al., 2011; Emerson et al., 2017).

Studies using stimulus-evoked fMRI in sleeping infants show that adult language regions such as the left superior temporal (STG) and inferior frontal (IFG) gyri are already engaged in language processing early in life (Dehaene-Lambertz et al., 2002; Perani et al., 2011; Wagner et al., 2025). While the brain’s language network features a robustly lateralized architecture in adulthood (Friederici & Gierhan, 2013), passive listening studies show that more bilateral language processing may be the norm at birth (Perani et al., 2011; Sato et al., 2012) and even to some extent in older infants (Wagner et al., 2025). Left-hemisphere (Shultz et al., 2014) and classical temporal language areas (Redcay et al., 2008) become increasingly selective for speech sounds throughout early life. Moreover, infants at high likelihood for ASD – a condition partially characterized by deficits in language and communication skills – show reduced experience-dependent recruitment of the left STG during statistical language learning (Liu et al., 2021) as well as diminished neural activation of left temporal language areas during speech processing (Wagner et al., 2025). This suggests that adult-like left-lateralization for language is a gradual maturational process that may follow an altered developmental course in conditions associated with language impairment.

Resting-state fMRI (rs-fMRI) studies show that interhemispheric connectivity for Broca’s and Wernicke’s areas is weak *in utero* (Thomason et al., 2013), but is robustly detected just after birth in 2-day-old neonates (Perani et al., 2011) and through the first month of life (Scheinost et al., 2022). By contrast, intrahemispheric connectivity between the inferior frontal and superior temporal nodes is immature during the perinatal period (Perani et al., 2011; Scheinost et al., 2022). Combined with evidence from longitudinal studies across longer periods of development (Emerson et al., 2016; Liu et al., 2023), the evidence suggests that core language regions begin to shift away from a symmetrical architecture and toward functional asymmetry after birth, with developmental curves that follow an inverted U-shape. Interestingly, one longitudinal rs-fMRI study in typically-developing infants showed that infants who follow a more pronounced quadratic trajectory of interhemispheric IFG connectivity display better language scores at 4-year follow-up (Emerson et al., 2016), indicating that a developmental trajectory of increasing symmetry in very early infancy, followed by decreasing symmetry, is favorable for frontal language regions. These later maturational reductions in interhemispheric connectivity of frontal and temporal language areas are also accompanied by increases in intrahemispheric connectivity, especially between core left-hemisphere areas (Emerson et al., 2016). Importantly, distinct functional modules across the brain follow regionally-specific developmental trajectories (Liu et al., 2023). Prior work examining longitudinal functional connectivity changes between nodes of the language network across the first year of life has demonstrated several maturational patterns that may lead to the adult-like configuration (Liu et al., 2020). These include increases in long-range connectivity (temporal to frontal, thalamus to frontal) and decreases in short-range connectivity (intra-temporal, intra-frontal, and thalamus to temporal). However, these results came from a relatively small sample (N = 20), and from an analysis that did not consider key cerebellar language regions.

In this study, we used rs-fMRI scans from the Baby Connectome Project (BCP; Howell et al., 2019) to investigate how the development of resting-sate functional connectivity within the infant language network relates to early receptive and expressive language development. Because rs-fMRI poses no demands on the participant, this approach is ideal for studying the development of functional brain connectivity in infants. In this well-powered, densely sampled cohort of typically-developing infants, *a priori* regions of the cerebrum (Friederici & Gierhan, 2013) and lobules of the cerebellum (Nettekoven et al., 2024) known to be involved in language processing were selected for a seed-based analysis of functional connectivity development to identify age-dependent maturational changes and associations with later language skills. These regions consisted of posterior superior and middle temporal gyri, the three subdivisions of the inferior frontal gyrus (pars orbitalis, triangularis, and opercularis), primary auditory cortex (Heschl’s gyrus), thalamus, as well as crus I and lobule VI of the cerebellum. Because language impairment affects around 10% of children (Norbury et al., 2016), we also used a data-driven machine learning approach to parse the behavioral heterogeneity in this sample. Participants were stratified into behavioral cohorts that had distinct trajectories of language development to examine how these might relate to language outcome.

## 2. Methods

### 2.1 Participants

Participants from this study are from the Baby Connectome Project (BCP; Howell et al., 2019), a longitudinal study of early brain and behavioral development in typically-developing infants. Data were accessed through the National Institute of Mental Health National Data Archive (Collection: UNC/UMN Baby Connectome Project). Eligible infants were born at 37 – 42 weeks gestational age, had an age-appropriate birth weight (>2000g), were free of major pregnancy and delivery complications, and had no family history of neurodevelopmental conditions such as ASD, intellectual disability, schizophrenia, or bipolar disorder. For the present study, we focus on resting-state functional scans collected within the first year of life (0 – 12 months), and behavioral assessments collected across the first three years of life (0 – 36 months).

Functional data from 314 MRI visits conducted from 0 – 12 months were considered for analysis. Of these, one was excluded for corrupted data, 38 for failing quality control during preprocessing, eight for excessive motion (mean framewise displacement > 0.40mm), and two for failing ICA-based motion correction. This left a final sample of 267 imaging visits from 154 unique participants retained for our analyses.

### 2.2 Behavioral Data + Behavioral Clustering

The Mullen Scales of Early Learning (MSEL; Mullen, 1995) were collected beginning at 3 months of age. This assessment yields age-normed t scores, age equivalent scores, and percentile rankings for five subdomains of development: gross motor, fine motor, receptive language, expressive language, and visual reception. The MacArthur-Bates Communicative Development Inventories (MCDI; Fenson, 2007) were administered from 12 – 24 months to index receptive and expressive language, as well as communicative gesture use. To assess theory of mind skills, which are closely intertwined with early language development (Ebert, 2020), the Children’s Social Understanding Scale (CSUS; Tahiroglu et al., 2014) was collected from 24 months onward. The Vineland Adaptive Behavior Scales (VABS II; Sparrow et al., 2005) were collected beginning at 3 months to assess child behavior in communication, socialization, daily living skills, and motor skills. The Dimensional Joint Attention Assessment (DJAA; Elison et al., 2013), collected from 8 – 16 months, assesses joint attention skills, which are necessary for advanced communication skills and interpersonal interactions. Behavioral assessment data were divided into age bins for group comparisons. See Figure S1 for a visualization of the age distributions for each measure.

As in a prior study (Okada et al., 2021), longitudinal k-means clustering (KmL package in R; Genolini et al., 2015) was used to partition participants into three cohorts based on developmental receptive language trajectories on the MSEL. Receptive language scores were used in favor of expressive language scores because receptive ability develops first (Kuhl, 2004), and shows greater early impairments in toddlers diagnosed with ASD (Hudry et al., 2010; Luyster et al., 2008). Because language assessments performed younger than 6 months of age tend to be less reliable due to infants’ limited behavioral repertoires, and because assessments were sparse past 30 months of age, language scores from 6 – 30 months of age were included in the clustering approach. Participants were included in the longitudinal clustering model if they had at least two MSEL assessments at different timepoints. Missing data points were estimated using KmL’s copy mean imputation (Genolini et al., 2013).

### 2.3 MRI acquisition

All MRI data were acquired during natural sleep on Siemens Prisma 3T scanners with 32-channel head coils, at the University of Minnesota and the University of North Carolina, Chapel Hill. A scout localizer, modified to be quieter than typical localizers, was acquired with a TR of 30ms and TE of 5ms. T2-weighted structural images were acquired with a 3D variable flip angle turbo spin-echo sequence with the following parameters: TR=3200ms, TE=564ms, matrix size=320×320, FOV=256mm, 208 sagittal slices, 0.8 x 0.8 x 0.8 mm^3^ resolution. Resting-state fMRI scans were collected as single-shot EPI sequences in two phase-encoding directions (anterior and posterior) to better correct for geometric distortions. These were collected with the following parameters: TR=800ms, TE=37ms, matrix size=104 x 91, FOV=208mm, 72 axial slices, 2 x 2 x 2 mm^3^ resolution. Resting-state scans lasted 5 min. 47s, and each participant ranged from one to four scans collected for a maximum amount of data of 23 min. 8s. Expanded details on image acquisition can be found in Howell et al. (2019).

Infant scans were conducted during natural sleep. Caregivers were encouraged to emulate their typical bedtime routine in preparation for the scan. Sound attenuating foam was used to line the bore of the scanner, and earplugs were placed in the infant’s ears and covered with tape for hearing protection. Sources of light were dimmed or turned off to aid the infant in falling asleep, and white noise was played in the background according to the caregiver’s preference. A trained member of the study staff remained in the scanner suite with the infant to monitor for signs of wakefulness or distress.

### 2.4 fMRI data preprocessing

Imaging data were preprocessed using FMRIB’s Software Library (FSL; Smith et al., 2004) version 6.0.7.4 unless otherwise stated. Structural scans underwent skull-stripping using Freesurfer’s (v7.4.1) SynthStrip tool, which uses deep learning trained on many image types (including infant data) to achieve optimal brain extraction. Functional scans underwent rigid-body motion correction in six directions before being co-registered to the infant’s own T2-weighted structural scan with six degrees of freedom (DOF) and then registered to a 1-year standard template derived from BCP data (Chen et al., 2022) with 12 DOF. Registration was manually inspected for quality assurance. Functional data were then smoothed using a 6-mm Gaussian kernel. Independent Component Analysis – Automatic Removal of Motion Artifacts (ICA-AROMA; Pruim et al., 2015) was used to identify and remove signal components attributable to motion or thermal, electrical, or physiological noise. This approach has been shown to improve reproducibility in resting-state analyses while preserving temporal DOF (Carone et al., 2017). Bandpass filtering (0.01 Hz < *t* < 0.1 Hz) was applied to further remove physiological noise such as heartbeat and respiration. Segmentation was performed on the structural scan using Freesurfer’s SynthSeg tool to derive masks of cerebrospinal fluid and white matter, which were then registered to the functional data in FSL. These masks were then used to regress mean white matter and CSF timeseries out of each functional run. Global signal regression (GSR) was also applied, as it has been shown to be effective for denoising when paired with ICA-AROMA (Parkes et al., 2018).

### 2.5 fMRI data analysis

A seed-based analytical approach was applied to examine functional connectivity between key nodes of the language network. Seeds were defined as cortical regions classically involved in language processing according to Friederici & Gierhan, 2013 (inferior frontal gyrus, IFG: pars orbitalis, triangularis, and opercularis; posterior superior temporal gyrus, pSTG – defined as the posterior third of the gyrus, as in Liu et al., 2020; and middle temporal gyrus, MTG) as well as the bilateral thalamus and lobules of the cerebellum implicated in language and social cognition (crus I and lobule VI; King et al., 2019).

For each participant, time series were extracted from these *a priori* language regions of interest (ROIs; left and right hemisphere separately, except for the thalamus) at the single-run level, and were subsequently correlated with the timeseries of every other voxel in the brain. The resultant correlation maps were then converted to z-statistic maps using Fisher’s r-to-z transformation. For participants with multiple resting-state scans, individual functional runs were combined in FSL FEAT to derive an average connectivity map for each participant.

In order to appropriately model connectivity in the full dataset of 267 scans, age analyses were conducted in R to model repeated observations. Pairwise correlations were computed for all ROI pairings, yielding a 17×17 connectivity matrix of eight bilateral ROIs, plus the thalamus. These pairwise relationships were then used as outcome variables in multilevel linear regression models to investigate changes in connectivity strength throughout the language network, modeled as a function of age. All models controlled for the effect of data collection site as a nuisance regressor. P-values were FDR corrected for 136 comparisons.

Group-level bottom-up analyses were also conducted to address three aims: 1) To assess the relationship between age and language network connectivity by using postnatal age as a bottom-up regressor in FSL (N = 154), 2) To examine the relationship between behavioral language measures and language connectivity across the entire group, using MSEL receptive and expressive language scores collected from 12-15 months as bottom-up behavioral regressors (N = 113), and 3) To compare language network connectivity between developmental language cohorts (as described above; N = 114). Because FSL cannot model repeated observations with missing data, each analysis used scans from unique infants who were selected to balance representation across the entire age range. Analyses 1 and 2 (described above) were masked by language regions (Figure S2), whereas analysis 3 examined group differences across the whole brain. Group-level analyses were performed in FSL FEAT using FMRIB’s Local Analysis of Mixed Effects (FLAME 1), with data collection site included as a demeaned nuisance regressor. Age was also included as a nuisance regressor when comparing connectivity across behavioral cohorts and when regressing connectivity data onto language scores. All group-level analyses were thresholded at Z > 3.1, cluster-corrected for multiple comparisons at P < 0.05. Significant cluster peaks are reported in Table 1. Demographic and MRI quality information for analyses 1 and 2 are reported in Table S1, and these same data for analysis 3 are reported in Table 2.

**Table 1.**
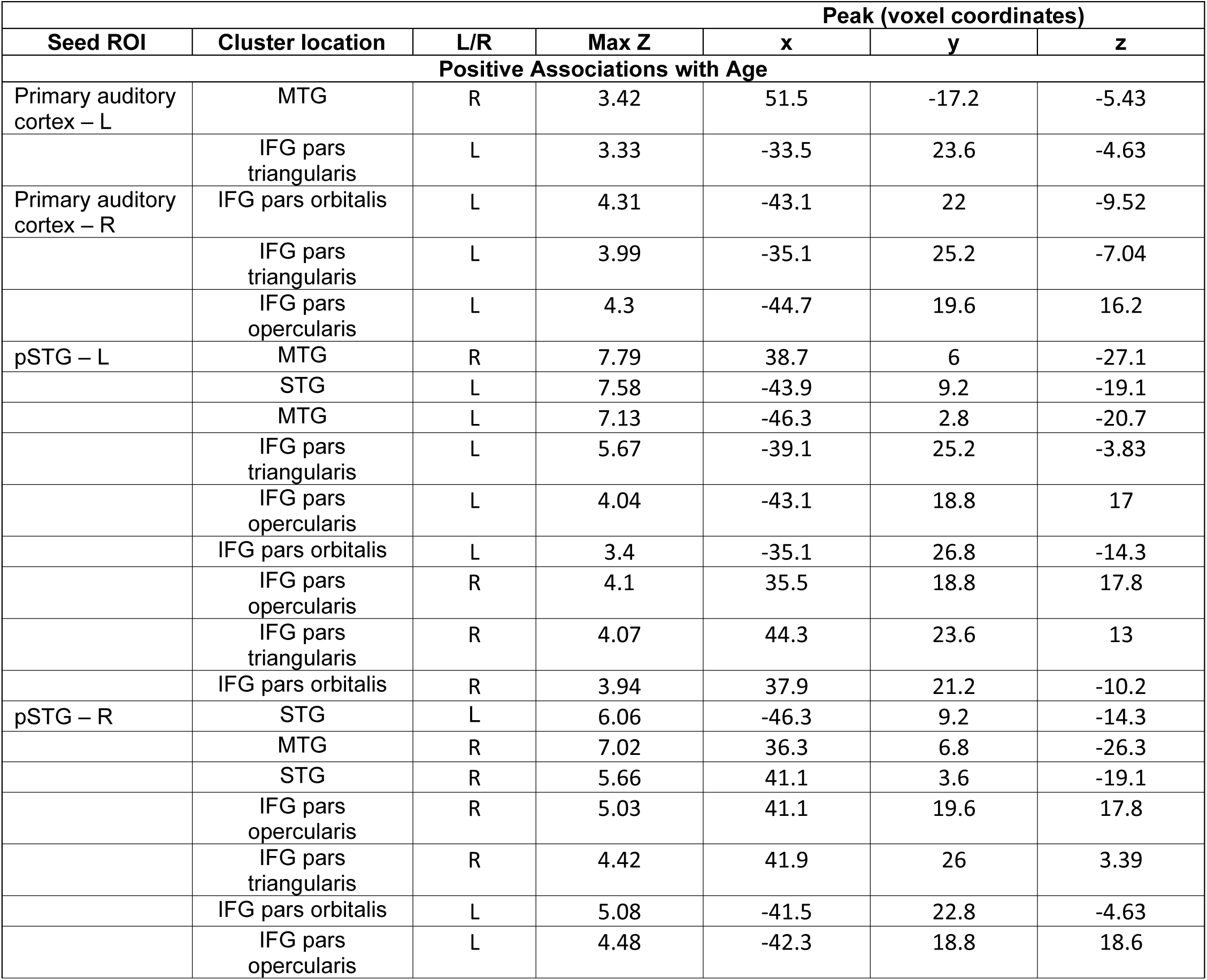
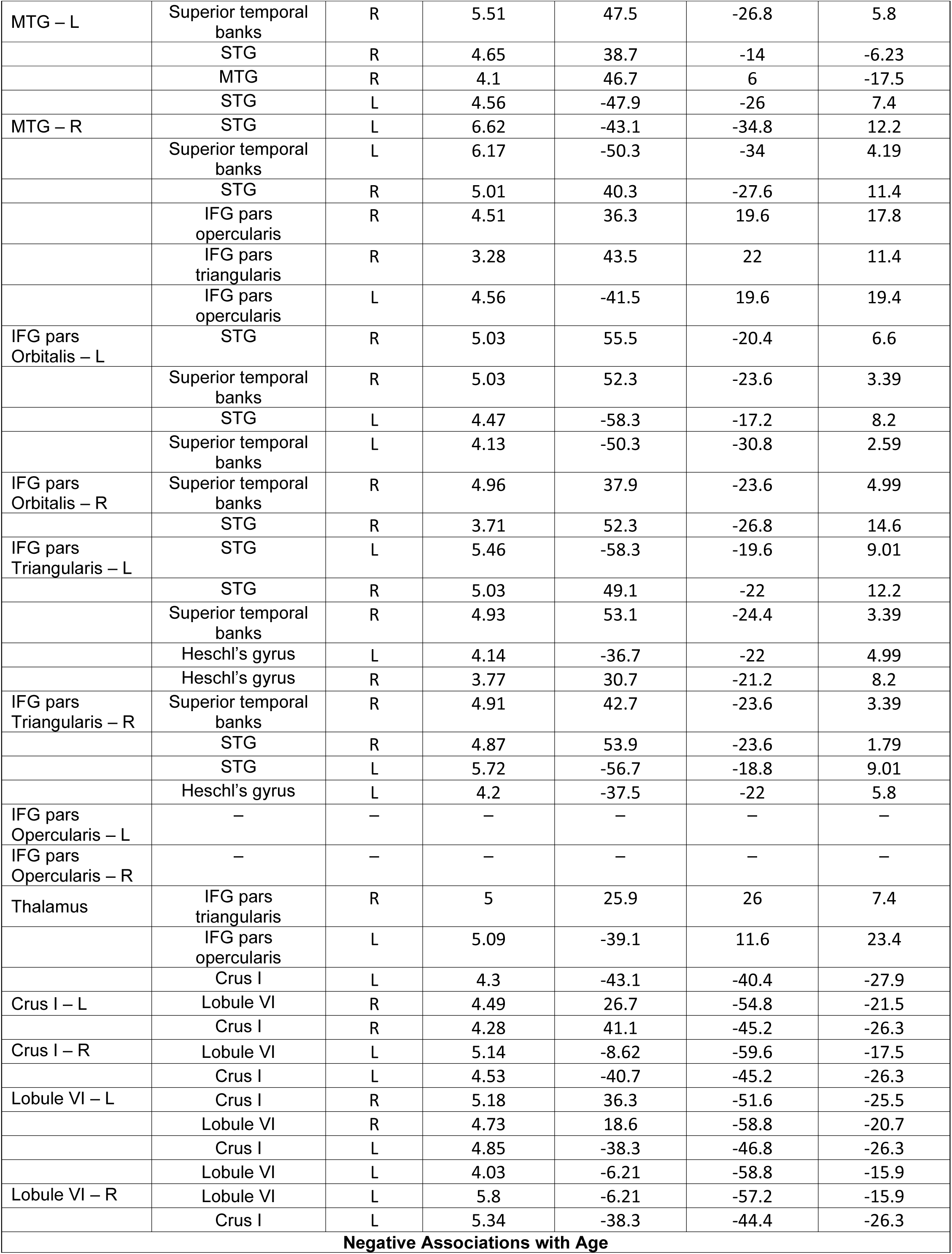
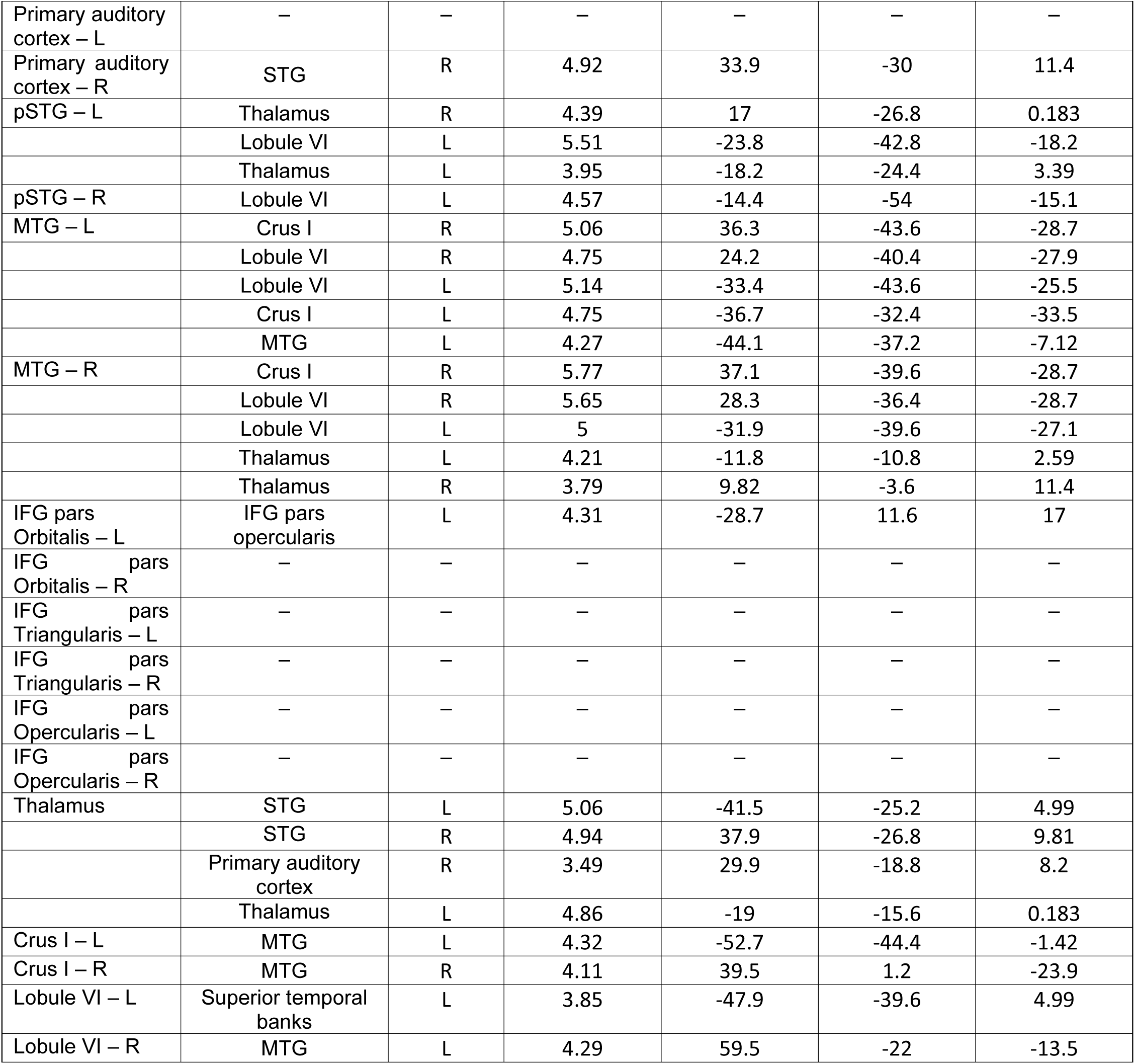
Coordinate table for language network connectivity positively and negatively associated with age across the subset of 154 scans from unique participants. STG: Superior temporal gyrus; pSTG: Posterior superior temporal gyrus; MTG: Middle temporal gyrus; IFG: Inferior frontal gyrus; L: Left; R: Right.

**Table 2.** Demographic information and behavioral assessments for the whole behavioral sample included in the clustering analysis, as well as the subset with neuroimaging data, subdivided by behavioral cohort. Abs= Absolute; Rel= Relative; UMN = University of Minnesota; UNC = University of North Carolina; ICA = Independent Component Analysis.

| Behaviorally Clustered Sample (N = 203) |  |  |  |  |  |
| --- | --- | --- | --- | --- | --- |
|  | Whole Sample: Mean or % | Advanced | Typical | Delayed | p (F or X<sup>2</sup>) |
| N | 203 | 65 | 113 | 25 | – |
| Sex |  |  |  |  | 0.44 |
| Female | 103 (51%) | 37 (57%) | 55 (49%) | 11 (44%) |  |
| Male | 100 (49%) | 28 (43%) | 58 (51%) | 14 (56%) |  |
| Ethnicity |  |  |  |  | **0.009\*\*** |
| Hispanic | 15 (7%) | 1 (2%) | 14 (12%) | 0 |  |
| Not Hispanic | 188 (93%) | 64 (98%) | 99 (88%) | 25 (100%) |  |
| Race |  |  |  |  | 0.17 |
| White | 156 (77%) | 55 (85%) | 87 (77%) | 14 (56%) |  |
| Black | 7 (3%) | 2 (3%) | 4 (4%) | 1 (4%) |  |
| Asian | 5 (2%) | 1 (1%) | 3 (3%) | 1 (4%) |  |
| More than one race | 35 (17%) | 7 (11%) | 19 (16%) | 9 (36%) |  |
| Site |  |  |  |  | 0.16 |
| UMN | 153 (75%) | 51 (78%) | 87 (77%) | 15 (60%) |  |
| UNC | 50 (25%) | 14 (22%) | 26 (23%) | 10 (40%) |  |
| Subset with Neuroimaging Data (N = 114 Unique Infants) |  |  |  |  |  |
|  | Whole Sample (N or mean) | Advanced | Typical | Delayed | p (F or X<sup>2</sup>) |
| N | 114 | 36 | 65 | 13 | – |
| Age at Scan | 6.7 ± 3.5 | 6.6 ± 3.5 | 6.6 ± 3.5 | 7.1 ± 3.5 | 0.92 |
| Sex |  |  |  |  | 0.36 |
| Female | 66 (58%) | 24 (67%) | 34 (52%) | 8 (61%) |  |
| Male | 48 (42%) | 12 (33%) | 31 (48%) | 5 (39%) |  |
| Ethnicity |  |  |  |  | **0.01\*** |
| Hispanic | 11 (10%) | 0 | 11 (17%) | 0 |  |
| Not Hispanic | 103 (90%) | 36 (100%) | 54 (83%) | 13 (100%) |  |
| Race |  |  |  |  | 0.26 |
| White | 87 (76%) | 28 (78%) | 51 (78%) | 8 (61%) |  |
| Black | 3 (3%) | 2 (6%) | 1 (2%) | 0 |  |
| Asian | 3 (3%) | 0 | 3 (5%) | 0 |  |
| More than one race | 21 (18%) | 6 (16%) | 10 (15%) | 5 (39%) |  |
| Site |  |  |  |  | 0.24 |
| UMN | 87 (76%) | 26 (72%) | 53 (82%) | 8 (61%) |  |
| UNC | 27 (24%) | 10 (28%) | 12 (18%) | 5 (39%) |  |
| Mean Abs. Motion | 0.34 ± 0.30 | 0.30 ± 0.21 | 0.34 ± 0.30 | 0.42 ± 0.49 | 0.46 |
| Mean Rel. Motion | 0.15 ± 0.06 | 0.15 ± 0.04 | 0.16 ± 0.06 | 0.15 ± 0.06 | 0.86 |
| Max Rel. Motion | 2.16 ± 2.42 | 1.84 ± 1.86 | 2.24 ± 2.76 | 2.60 ± 1.98 | 0.57 |
| # ICA Components Removed | 49.4 ± 18.6 | 49.1 ± 16.3 | 50.6 ± 19.7 | 44.2 ± 19.1 | 0.54 |

## 3. Results

Age-related changes in language network functional connectivity (FC) were modeled linearly using pairwise ROI-based multilevel models as well as seed-based regressions, yielding both a connectivity matrix of age-related changes in the language network as well as connectivity maps showing changes across the whole network.

### 3.1 ROI Matrix of Age-Dependent Functional Connectivity Changes

Overall, the connectivity matrix shows general patterns of increasing functional synchrony and specialization throughout the language network (Figure 1). Across the first year of life, thalamic connectivity significantly strengthened with several inferior frontal areas (bilateral pars opercularis and right pars triangularis), whereas thalamic connectivity with all temporal ROIs significantly decreased. Both left and right primary auditory cortices showed significant strengthening with all IFG subregions, with additional ipsilateral decreases in connectivity with the pSTG. Both left and right pSTG also showed widespread increases in connectivity strength with the IFG, with the exception of left pSTG to bilateral pars opercularis, and right pSTG to right pars opercularis. The bilateral pSTG also showed strong connectivity increases with bilateral MTG. Interestingly, out of all ROIs tested, the MTG (both left and right) was the only region to show reduced FC with cerebellar lobules as a function of advancing age. All cerebellar lobules (bilateral crus I and lobule VI) showed significant intra-cerebellar strengthening, with the exception of FC between left lobule VI and left crus I. Notably, bilateral lobule VI also showed strengthening with the left pars opercularis.

**Figure 1.**
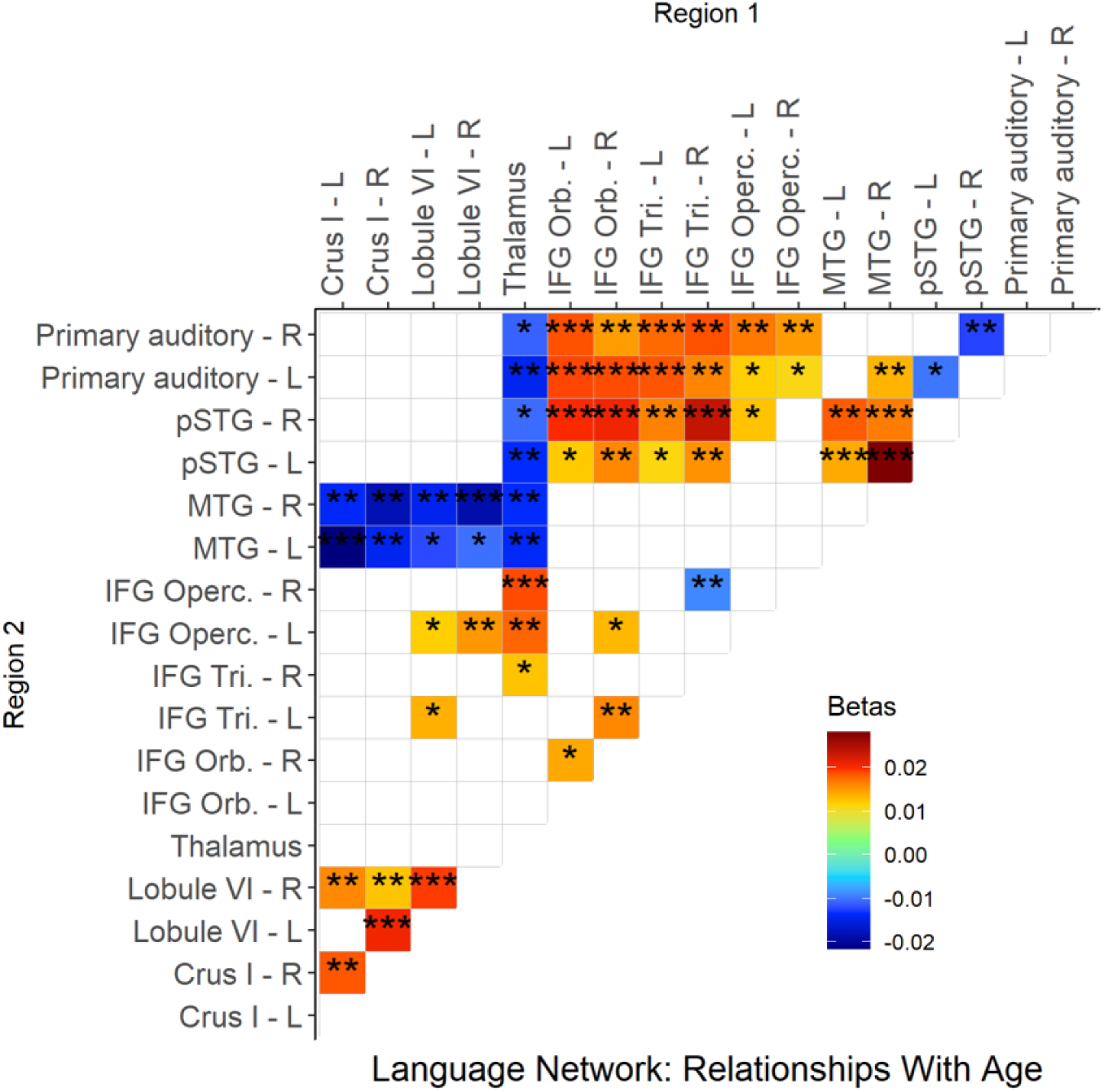
Changes in pairwise functional connectivity within language network ROIs as a function of age (0 – 12 months). Warm colors indicate linear increases with advancing age, while cool colors indicate linear reductions with advancing age. Plotted betas control for the effects of repeated measures and data collection site. Significance thresholds are FDR-corrected for multiple comparisons. *=*p*<0.05, **=*p*<0.01, ***=*p*<0.001. R: right; L: left; pSTG: posterior superior temporal gyrus; MTG: middle temporal gyrus; IFG: inferior frontal gyrus; Operc: opercularis; Tri: triangularis; Orb: orbitalis.

### 3.2 Bottom-Up Age Regressions

Bottom-up age regressions within the language network (Figures 2A-C; peaks reported in Table 1) provided a more detailed analysis of maturational relationships. Because FSL cannot model repeated observations with missing data, this analysis was carried out in a subset of 154 scans from unique infants. Overall, these results largely recapitulated those presented in Figure 1, with some exceptions, as detailed below, for each ROI.

**Figure 2A.**
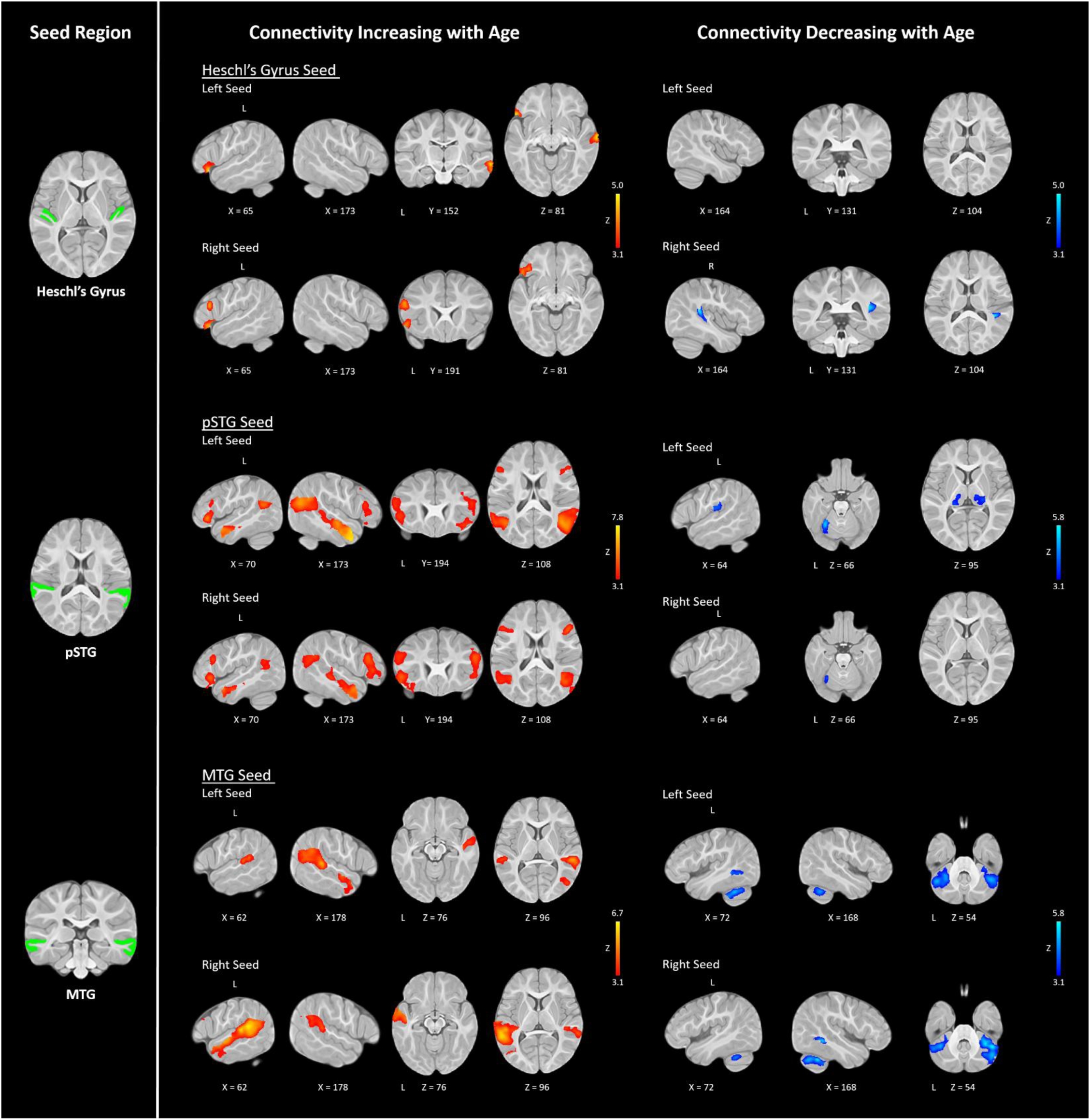
Temporal ROI Age Associations. Age-dependent changes in language network connectivity (N=154) for Heschl’s gyrus (primary auditory cortex), posterior superior temporal gyrus (pSTG), and middle temporal gyrus (MTG). Connectivity increases are shown in red colors, whereas connectivity decreases are shown in blue. Results were thresholded at Z>3.1. Seed ROIs are displayed in the left panel.

**Figure 2B.**
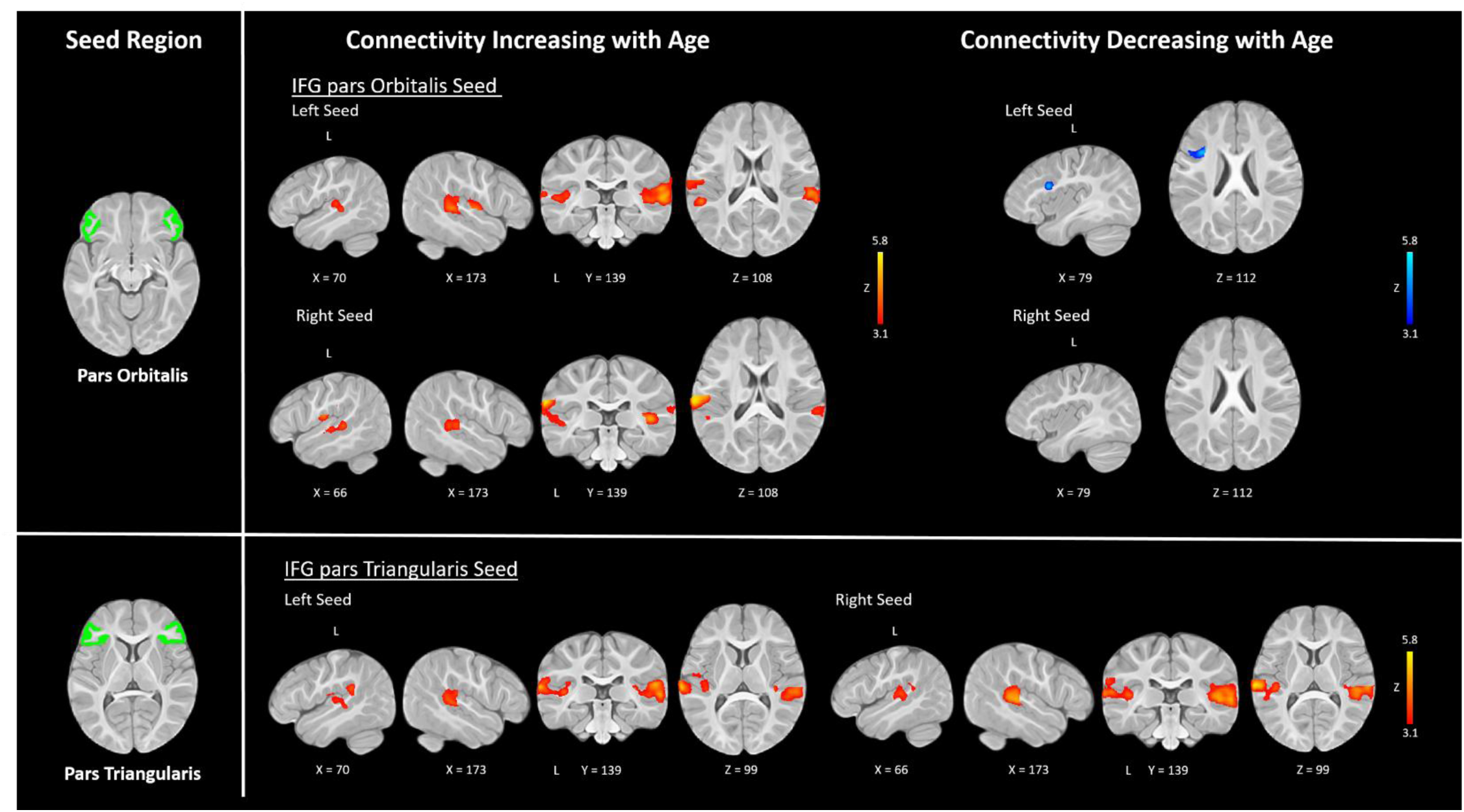
Frontal ROI Age Associations. Age-dependent changes in language network connectivity (N=154) for the pars orbitalis and pars triangularis of the inferior frontal gyrus (IFG). No significant associations were found for the pars opercularis. Connectivity increases are shown in red colors, whereas connectivity decreases are shown in blue. Results were thresholded at Z>3.1. Seed ROIs are displayed in the left panel.

**Figure 2C.**
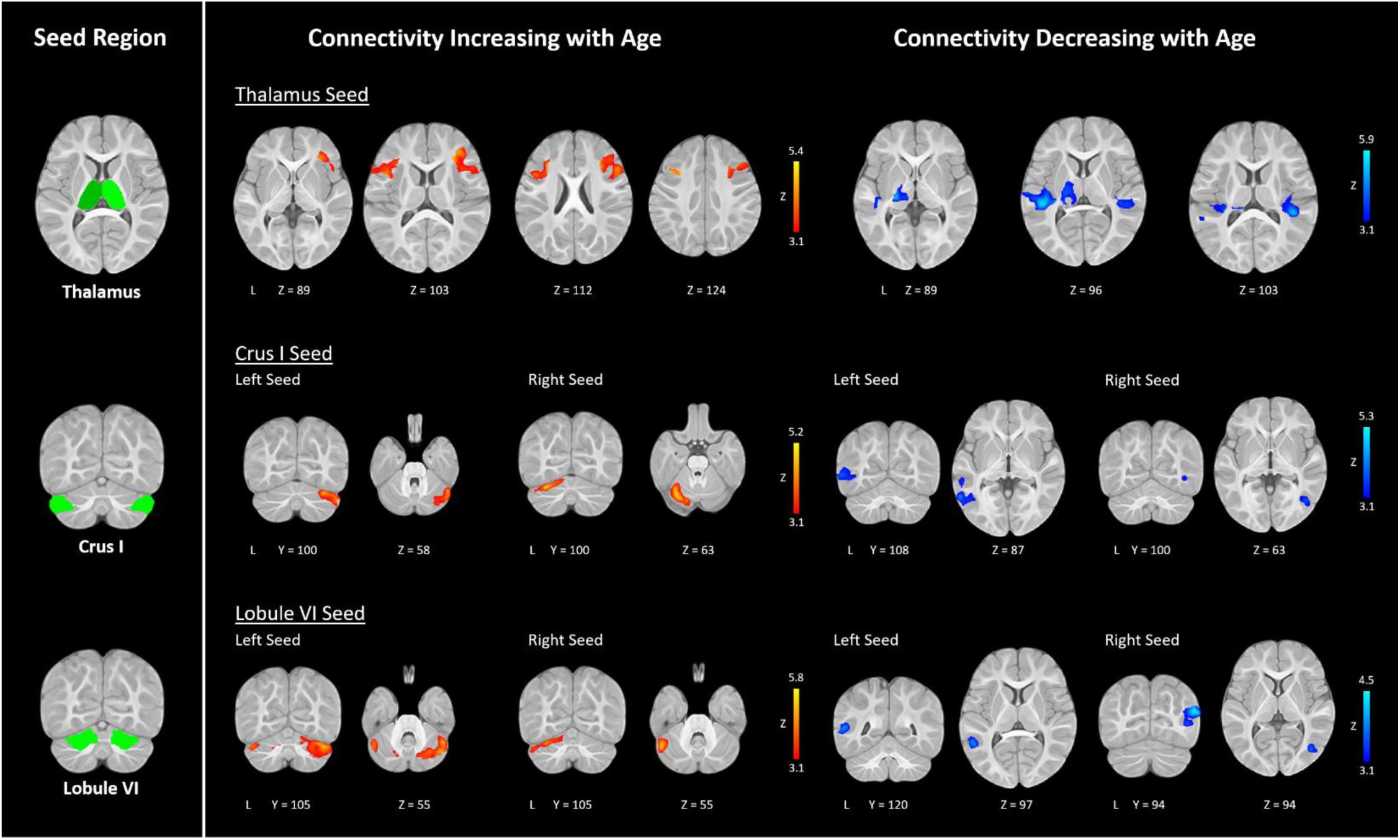
Subcortical ROI Age Associations. Age-dependent changes in language network connectivity (N=154) for the thalamus (bilateral), crus I, and lobule VI. Connectivity increases are shown in red colors, whereas connectivity decreases are shown in blue. Results were thresholded at Z>3.1. Seed ROIs are displayed in the left panel.

While the correlation matrix detected age-dependent FC increases between both primary auditory seeds and all subregions of the IFG, in the bottom-up analyses left primary auditory cortex had significant strengthening with ipsilateral pars triangularis as well as the right MTG (also found in Figure 1). Right primary auditory cortex showed significant strengthening with all IFG subregions, but only those in the left hemisphere. Whereas the pairwise ROI approach detected age-dependent FC decreases between primary auditory cortex and ipsilateral pSTG, in the bottom-up approach we found that only right primary auditory cortex underwent age-dependent decreases with an ipsilateral superior temporal cluster (Figure 2A).

Connectivity increases for the pSTG largely mirrored the ROI pairwise findings such that, on the whole, FC with MTG and subregions of the IFG underwent age-dependent strengthening. While the ROI pairwise analyses showed connectivity increases with bilateral pars orbitalis, pars triangularis, and MTG for the left pSTG, in the bottom-up regression the left pSTG also showed additional connectivity increases with the pars opercularis. Previously, the right pSTG showed increases with bilateral MTG and all IFG subregions except the right pars opercularis. However, in the bottom-up analysis there were age-dependent increases only in ipsilateral MTG connectivity, as well as in connectivity with left pars orbitalis, right pars triangularis, and bilateral pars opercularis. Both pSTG seeds showed decreasing FC with the thalamus in the ROI pairwise analyses, but in the whole-brain approach this relationship was only significant for the left pSTG seed. Additionally, bilateral pSTG also showed significant age-dependent decreases in FC with left cerebellar lobule VI in the bottom-up analyses (Figure 2A).

MTG connectivity with bilateral superior temporal areas strengthened with advancing age, mirroring the ROI pairwise results as well as the pSTG seed results. We also found that the right MTG showed additional connectivity increases in the bottom-up analyses with multiple IFG subregions, including the bilateral pars opercularis and the right pars triangularis. This analysis also reflected the strong age-dependent decreases in connectivity previously observed between the MTG and cerebellar ROIs, as well as the thalamus. The left MTG showed age-dependent decreases in connectivity with all cerebellar ROIs, while the right MTG showed decreases for bilateral lobule VI, right crus I, and bilateral thalamus (Figure 2A).

While in our ROI pairwise analyses the pars orbitalis and triangularis showed bilateral connectivity increases with bilateral superior temporal cortices, in the bottom-up analyses we detected only ipsilateral strengthening between right pars orbitalis and superior temporal areas. The pars triangularis also showed increases in connectivity with primary auditory cortex, with the left seed showing increases with bilateral auditory cortex, and the right seed showing only contralateral connectivity increases. The pars opercularis seeds, however, showed no statistically significant connectivity increases with advancing age. An age-dependent connectivity decrease was detected for the left pars orbitalis seed, which showed decreasing connectivity with a cluster in the ipsilateral pars opercularis (Figure 2B).

Similar to the ROI pairwise analyses, the thalamus showed age-related increases in connectivity with the right pars triangularis and left pars opercularis. There was also a significant positive cluster in the left crus I. In line with the broad reductions in thalamic-temporal connectivity observed in the prior analyses, age-related decreases were found in relation to bilateral STG and right primary auditory cortex (Figure 2C).

As in the ROI pairwise analyses, intracerebellar connectivity underwent significant strengthening with increasing age. In the bottom-up analyses, left lobule VI underwent significant increases in connectivity with every other cerebellar ROI, while right lobule VI, left crus I, and right crus I showed increases with contralateral cerebellar ROIs. Although we did not see such consistent FC decreases between cerebellar ROIs and bilateral MTG as in the ROI pairwise analysis, we did observe age-dependent FC decreases between crus I seeds and the ipsilateral MTG, as well as a decrease in connectivity between right lobule VI and left MTG. Left lobule VI also showed FC decreases with ipsilateral superior temporal cortex (Figure 2C).

### 3.3 Connectivity Relationships with MSEL Language Measures

Two ROIs – right primary auditory cortex and right crus I – showed functional connectivity with other language regions that was significantly associated with later receptive language scores. Better receptive language scores at 12 – 15 months were predicted by stronger interhemispheric connectivity between right primary auditory cortex and a left middle and inferior temporal cluster (Figure 3A). Conversely, interhemispheric cerebellar connectivity appeared to be detrimental to language development, with stronger connectivity between the right crus I and left cerebellar ROIs (left crus I and lobule VI) predicting worse receptive language scores (Figure 3B). There were no significant functional connectivity relationships with expressive language scores.

**Figure 3.**
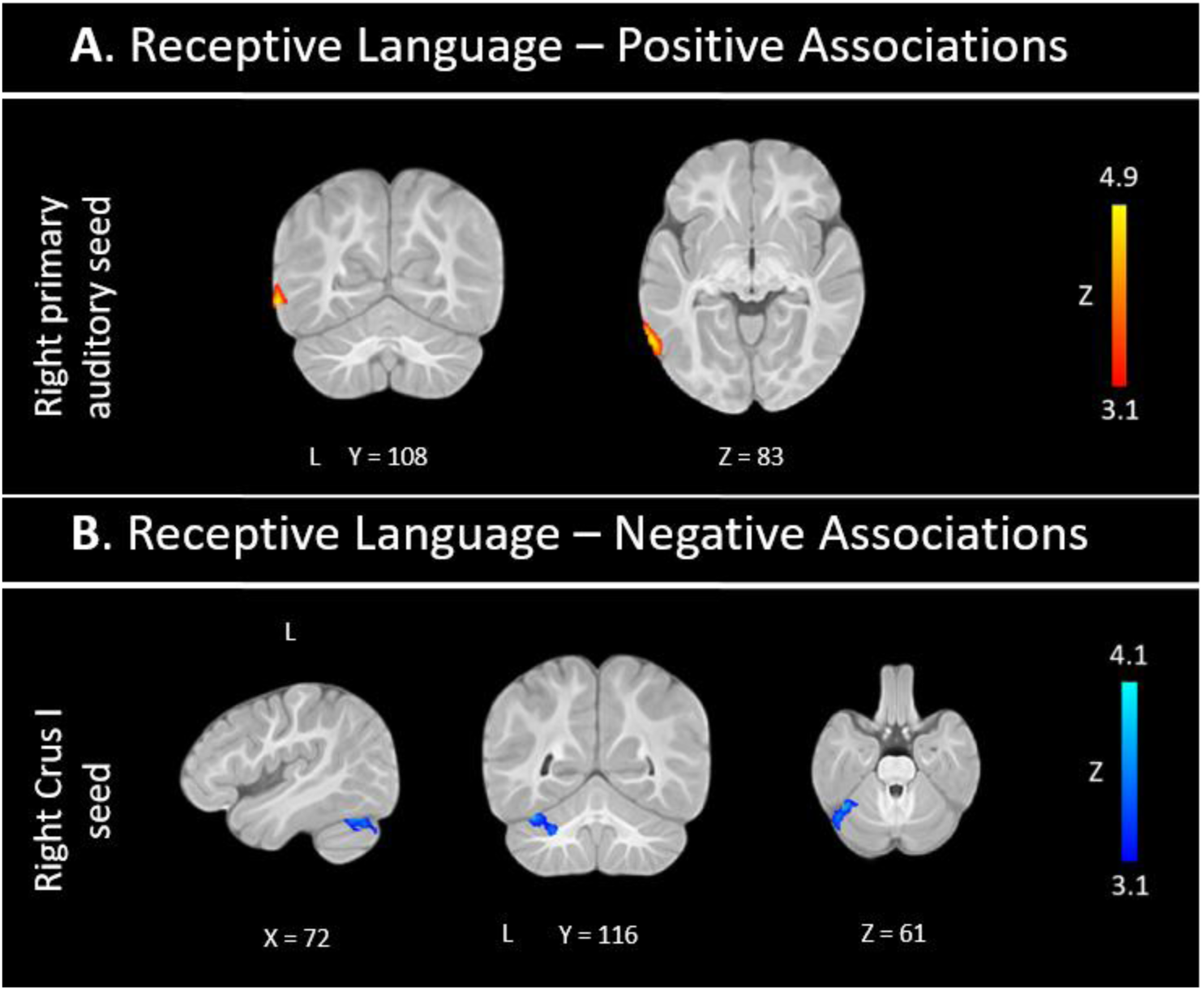
Brain associations with the Mullen Receptive Language subscale at approximately 12 months of age (N=113). These regressions were masked by language network ROIs. Warm colors indicate connectivity positively associated with receptive language skills (A), while cool colors indicate connectivity negatively associated with receptive language skills (B).

### 3.4 Behavioral Clustering

A total of 203 infants had sufficient behavioral data (i.e., at least two observations) for longitudinal k-means clustering. This sample was partitioned into three groups: 55.7% (N=113) of the infants were categorized as having a “Typical” language development trajectory; 32% (N=65) were categorized as “Advanced”; and 12.3% (N=25) were categorized as “Delayed”. Group trajectories on the receptive (Figure 4A) and expressive (Figure 4B) language subscales are displayed, with boxplots illustrating the differences across the three groups in Figure S3. These cohorts did not significantly differ on sex, data collection site, or race, but they were unmatched on ethnicity (Table 2).

**Figure 4.**
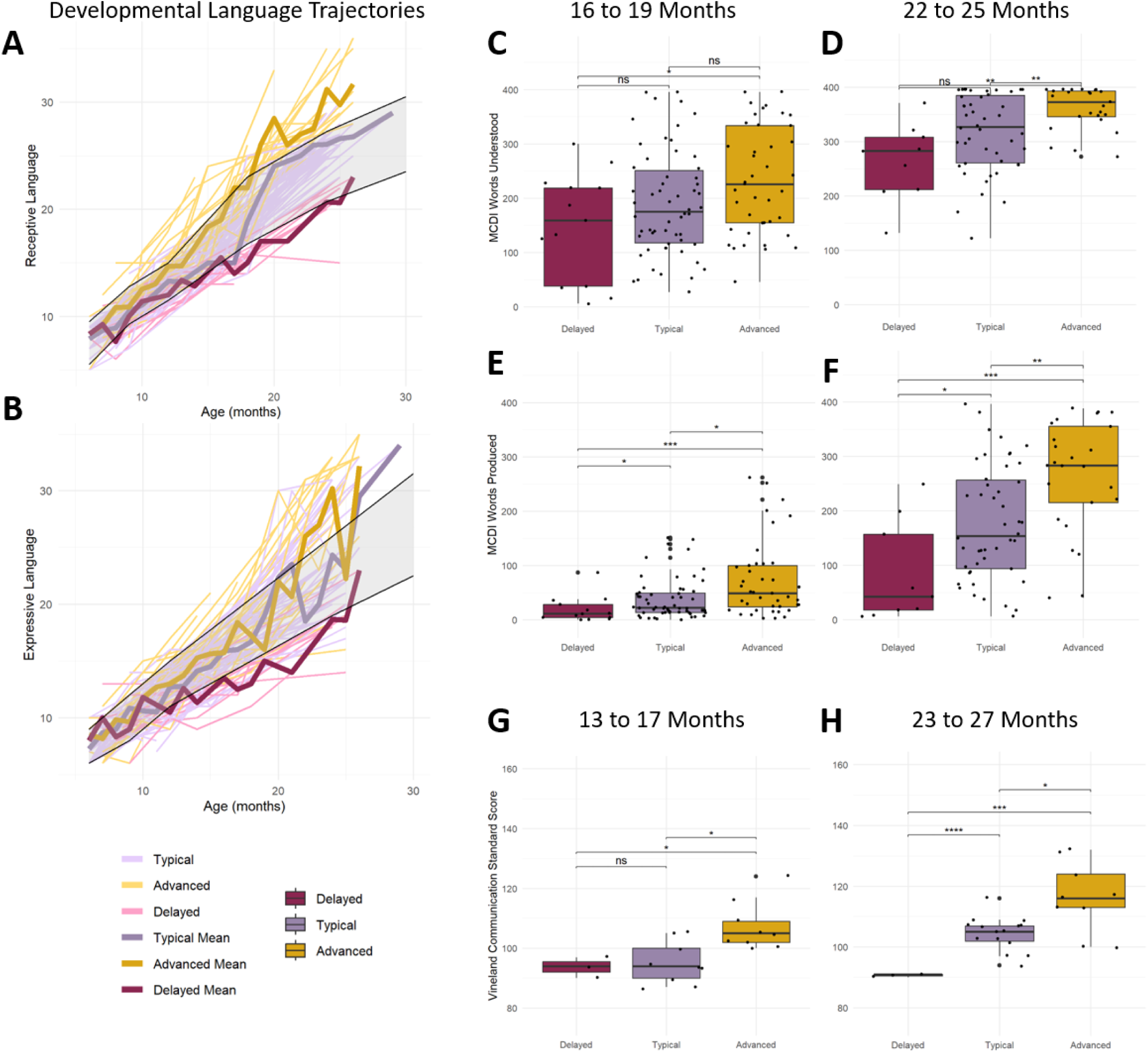
Cohort differences on language skills. Developmental language trajectories on the MSEL receptive language (A) and expressive language (B) subscales. Although age-normed scores were used in the clustering analysis, these trajectories present raw scores for visualization purposes. The normative range is shaded in gray. The cohorts differed on the number of words understood according to the MCDI at 16 to 19 months (C; F(2) = 4.44, *p* = 0.014) and 22 to 25 months (D; F(2) = 8.54, *p* < 0.001), as well as on number of MCDI words produced at these same ages (E; F(2) = 7.62, *p* < 0.001) and (F; F(2) = 11.52, *p* < 0.001). The cohorts also differed on the Vineland communication subscale at 13 to 17 months (G; F(2) = 7.45, *p* = 0.004) and 23 to 27 months (H; F(2) = 13.17, *p* < 0.001). \**p*<0.05, \*\**p*<0.01, \*\*\**p*<0.001, \*\*\*\**p*<0.0001. MSEL = Mullen Scales of Early Learning; MCDI = MacArthur-Bates Communicative Development Inventories.

A full summary of group differences (ANOVA statistics and pairwise group comparisons) on all behavioral assessments across various age bins is provided in Table 3. Not surprisingly, the three groups significantly differed on MSEL receptive language T-scores – the subscale from which the group partitions were derived – beginning at 9 months of age. The groups also significantly differed on expressive language trajectories beginning at 12 months of age. With the exception of MSEL receptive language scores at 9-11 months, wherever the three groups significantly differed according to ANOVAs, the Advanced group consistently scored higher than the Typical group, who in turn scored higher than the Delayed group, on all behavioral assessments.

**Table 3.** Behavioral scores for developmental assessments across developmental language cohorts. Mean ± standard deviation reported for each group. Significance levels for pairwise two-sample t-tests are shown with asterisks: * *p*<0.05, ** *p*<0.01, \*\*\**p*<0.001, \*\*\*\**p*<0.0001. F- and p-values reflect the results of one-way ANOVAs across the three groups. Adv = Advanced, T = Typical, D = Delayed. MSEL = Mullen Scales of Early Learning; MCDI = MacArthur Bates Communicative Development Inventories; CSUS = Children’s Social Understanding Scale; DJAA = Dimensional Joint Attention Assessment; VABS = Vineland Adaptive Behavior Scales.

| Age Bin | Advanced | Typical | Delayed | Adv v. T | Adv v. D | T v. D | F | p |
| --- | --- | --- | --- | --- | --- | --- | --- | --- |
| MSEL Receptive Language T-scores |  |  |  |  |  |  |  |  |
| 6–8 mo. | 54.1 ± 9.79 | 52.0 ± 8.51 | 49.1 ± 11.1 | — | — | — | 1.11 | 0.34 |
| 9–11 mo. | 53.5 ± 8.02 | 47.2 ± 7.78 | 48.4 ± 10.7 | ** | — | — | 5.48 | **0.006** |
| 12–15 mo. | 52.9 ± 9.77 | 44.9 ± 6.65 | 42.2 ± 8.98 | **** | ** | — | 16.9 | **<0.001** |
| 16–19 mo. | 58.8 ± 9.52 | 44.0 ± 11.7 | 36.4 ± 5.40 | *** | **** | * | 15.6 | **<0.001** |
| 22–25 mo. | 67.8 ± 7.08 | 55.9 ± 4.38 | 38.9 ± 6.75 | **** | **** | *** | 84.2 | **<0.001** |
| MSEL Expressive Language T-scores |  |  |  |  |  |  |  |  |
| 6–8 mo. | 53.8 ± 9.77 | 52.9 ± 7.80 | 54.9 ± 10.3 | — | — | — | 0.27 | 0.76 |
| 9–11 mo. | 54.0 ± 9.51 | 49.6 ± 9.74 | 53.4 ± 11.0 | — | — | — | 2.02 | 0.14 |
| 12–15 mo. | 53.3 ± 8.04 | 49.5 ± 8.89 | 42.9 ± 7.99 | * | ** | * | 8.55 | **<0.001** |
| 16–19 mo. | 49.9 ± 8.96 | 48.7 ± 5.09 | 38.5 ± 5.55 | — | ** | ** | 8.11 | **<0.001** |
| 22–25 mo. | 61.7 ± 13.0 | 51.8 ± 11.6 | 39.0 ± 5.18 | ** | **** | **** | 11.9 | **<0.001** |
| MSEL Visual Reception T-scores |  |  |  |  |  |  |  |  |
| 6–8 mo. | 58.3 ± 8.91 | 57.5 ± 10.2 | 55.4 ± 10.9 | — | — | — | 0.30 | 0.74 |
| 9–11 mo. | 55.8 ± 11.0 | 52.7 ± 8.65 | 51.1 ± 10.0 | — | — | — | 1.22 | 0.30 |
| 12–15 mo. | 50.7 ± 7.94 | 51.5 ± 9.19 | 52.5 ± 8.92 | — | — | — | 1.26 | 0.77 |
| 16–19 mo. | 48.3 ± 12.1 | 46.6 ± 10.5 | 48.6 ± 6.39 | — | — | — | 0.16 | 0.85 |
| 22–25 mo. | 68.4 ± 8.27 | 58.7 ± 8.28 | 48.4 ± 13.4 | *** | ** | — | 16.25 | **<0.001** |
| MSEL Fine Motor T-scores |  |  |  |  |  |  |  |  |
| 6–8 mo. | 53.2 ± 12.3 | 51.3 ± 10.2 | 51.2 ± 9.43 | — | — | — | 0.25 | 0.78 |
| 9–11 mo. | 57.8 ± 10.7 | 55.7 ± 10.3 | 58.9 ± 6.89 | — | — | — | 1.22 | 0.30 |
| 12–15 mo. | 59.5 ± 8.47 | 59.5 ± 8.64 | 59.0 ± 7.14 | — | — | — | 0.02 | 0.98 |
| 16–19 mo. | 52.1 ± 4.81 | 52.7 ± 8.37 | 57.1 ± 6.94 | — | — | — | 1.47 | 0.24 |
| 22–25 mo. | 56.4 ± 7.62 | 52.5 ± 9.82 | 45.5 ± 10.6 | — | — | — | 3.88 | **0.03** |
| MSEL Gross Motor T-scores |  |  |  |  |  |  |  |  |
| 6–8 mo. | 51.0 ± 8.20 | 52.3 ± 9.01 | 53.6 ± 9.78 | — | — | — | 0.35 | 0.71 |
| 9–11 mo. | 50.8 ± 6.41 | 48.6 ± 8.16 | 49.7 ± 9.11 | — | — | — | 0.78 | 0.46 |
| 12–15 mo. | 52.5 ± 9.73 | 51.1 ± 11.5 | 45.6 ± 13.6 | — | — | — | 2.07 | 0.13 |
| 16–19 mo. | 53.9 ± 7.27 | 52.3 ± 8.25 | 39.5 ± 13.0 | — | — | — | 7.42 | **0.002** |
| 22–25 mo. | 53.4 ± 14.4 | 47.8 ± 9.10 | 43.4 ± 10.9 | — | — | — | 2.96 | **0.06** |
| MCDI Words Comprehended |  |  |  |  |  |  |  |  |
| 12–15 mo. | 103.42 ± 83.79 | 85.65 ± 72.19 | 69.00 ± 56.84 | — | — | — | 1.07 | 0.347 |
| 16–19 mo. | 233.34 ± 98.80 | 190.41 ± 96.99 | 148.54 ± 99.31 | * | * | — | 4.44 | **0.014** |
| 22–25 mo. | 361.08 ± 38.92 | 317.80 ± 71.61 | 264.11 ± 71.42 | ** | ** | — | 8.54 | **<0.001** |
| MCDI Words Produced |  |  |  |  |  |  |  |  |
| 12–15 mo. | 11.82 ± 17.92 | 7.48 ± 7.08 | 3.64 ± 4.39 | — | * | * | 2.62 | 0.078 |
| 16–19 mo. | 72.00 ± 70.03 | 38.45 ± 37.79 | 19.00 ± 24.30 | ** | *** | * | 7.62 | **<0.001** |
| 22–25 mo. | 264.80 ± 102.98 | 177.76 ± 105.34 | 84.11 ± 92.55 | ** | *** | * | 11.52 | **<0.001** |
| CSUS Score |  |  |  |  |  |  |  |  |
| 24–28 mo. | 1.88 ± 0.47 | 1.88 ± 0.30 | 1.77 ± 0.18 | — | — | — | 0.48 | 0.623 |
| 31–35 mo. | 2.71 ± 0.62 | 2.31 ± 0.49 | 2.27 ± 0.48 | ** | ** | — | 6.34 | **0.003** |
| DJAA Score |  |  |  |  |  |  |  |  |
| 8–11 mo. | 7.50 ± 3.65 | 6.98 ± 3.88 | 8.38 ± 4.96 | — | — | — | 0.236 | 0.79 |
| 12–16 mo. | 11.33 ± 2.88 | 11.23 ± 3.92 | 12.10 ± 5.27 | — | — | — | 0.105 | 0.90 |
| VABS Communication Standard Score |  |  |  |  |  |  |  |  |
| 8–12 mo. | 111.67 ± 9.52 | 98.83 ± 17.48 | 95.50 ± 19.09 | — | — | — | 1.55 | 0.24 |
| 13–17 mo. | 107.11 ± 8.28 | 95.11 ± 6.90 | 93.67 ± 3.51 | ** | ** | — | 7.45 | **0.004** |
| 23–27 mo. | 116.22 ± 11.60 | 104.47 ± 5.59 | 90.67 ± 0.58 | * | *** | *** | 13.17 | **<0.001** |
| Overall 6–32 mo. | 111.31 ± 10.54 | 99.82 ± 12.52 | 93.00 ± 7.75 | *** | *** | — | 12.33 | **<0.001** |
| **VABS Socialization Standard Score** |  |  |  |  |  |  |  |  |
| 8–12 mo. | 111.67 ± 7.84 | 105.17 ± 6.06 | 101.50 ± 10.61 | — | — | — | 2.38 | 0.12 |
| 13–17 mo. | 110.33 ± 6.91 | 103.89 ± 6.37 | 103.00 ± 10.54 | — | — | — | 2.23 | 0.14 |
| 23–27 mo. | 113.44 ± 8.31 | 108.80 ± 6.09 | 100.00 ± 6.24 | — | * | — | 4.36 | **0.02** |
| Overall 6–32 mo. | 111.14 ± 7.55 | 106.32 ± 6.16 | 101.50 ± 7.80 | ** | ** | — | 7.82 | **<0.001** |
| **VABS Motor Skills Standard Score** |  |  |  |  |  |  |  |  |
| 8–12 mo. | 104.00 ± 12.17 | 99.25 ± 16.81 | 81.5 ± 2.12 | — | ** | ** | 1.68 | 0.22 |
| 13–17 mo. | 105.78 ± 3.77 | 100.44 ± 11.83 | 94.33 ± 11.59 | — | — | — | 1.95 | 0.17 |
| 23–27 mo. | 104.11 ± 7.17 | 102.00 ± 6.86 | 99.00 ± 3.61 | — | — | — | 0.70 | 0.51 |
| Overall 6–32 mo. | 104.76 ± 7.06 | 99.07 ± 13.79 | 92.88 ± 9.83 | * | * | — | 4.10 | **0.02** |

This partition of developmental language cohorts was further validated on independent language measures (i.e., the MCDI and VABS). The three groups significantly differed on MCDI receptive and productive vocabulary size from 16 months onward (Figure 4C-F), on the VABS Communication subscale at 13-17 and 23-27 months (Figure 4G, 4H), and the VABS Socialization subscale at 23-27 months. Notably, theory of mind abilities – closely related to language development – on the CSUS assessment also significantly differed across the three groups at 31-35 months. There were no group differences on the Dimensional Joint Attention Assessment at any age. Although they were clustered on a measure of receptive language, these three cohorts also differed on several non-language metrics. These included the MSEL Visual Reception and Fine Motor subscales at 22-25 months, the MSEL Gross Motor subscale at 16-19 months (trending at 22-25 months), and the VABS motor subscale across the whole age range of 6-32 months.

### 3.5 Functional Connectivity Differences across Developmental Language Cohorts

Out of the 203 participants who were assigned to developmental language cohorts, neuroimaging data were available for 114 infants. Within this subsample, the three groups did not significantly differ on age at the time of MRI scan, sex, race, data collection site, or MRI quality metrics (Table 2). However, the groups were unmatched on ethnicity such that all Hispanic participants (N=11) fell into the Typical group. When we examined group differences in functional connectivity between the three language cohorts for each language ROI, we identified several connectivity patterns with non-language regions that were associated with delayed language trajectories (peaks reported in Table 4). In post-hoc analyses, we also evaluated how these differences in connectivity might relate to later language measures. Since the receptive language measure was used to derive the three behavioral cohorts, these post-hoc analyses focused on the other available language measures (i.e., the MSEL expressive language subscale, the MCDI, and the VABS Communication subscale).

**Table 4.**
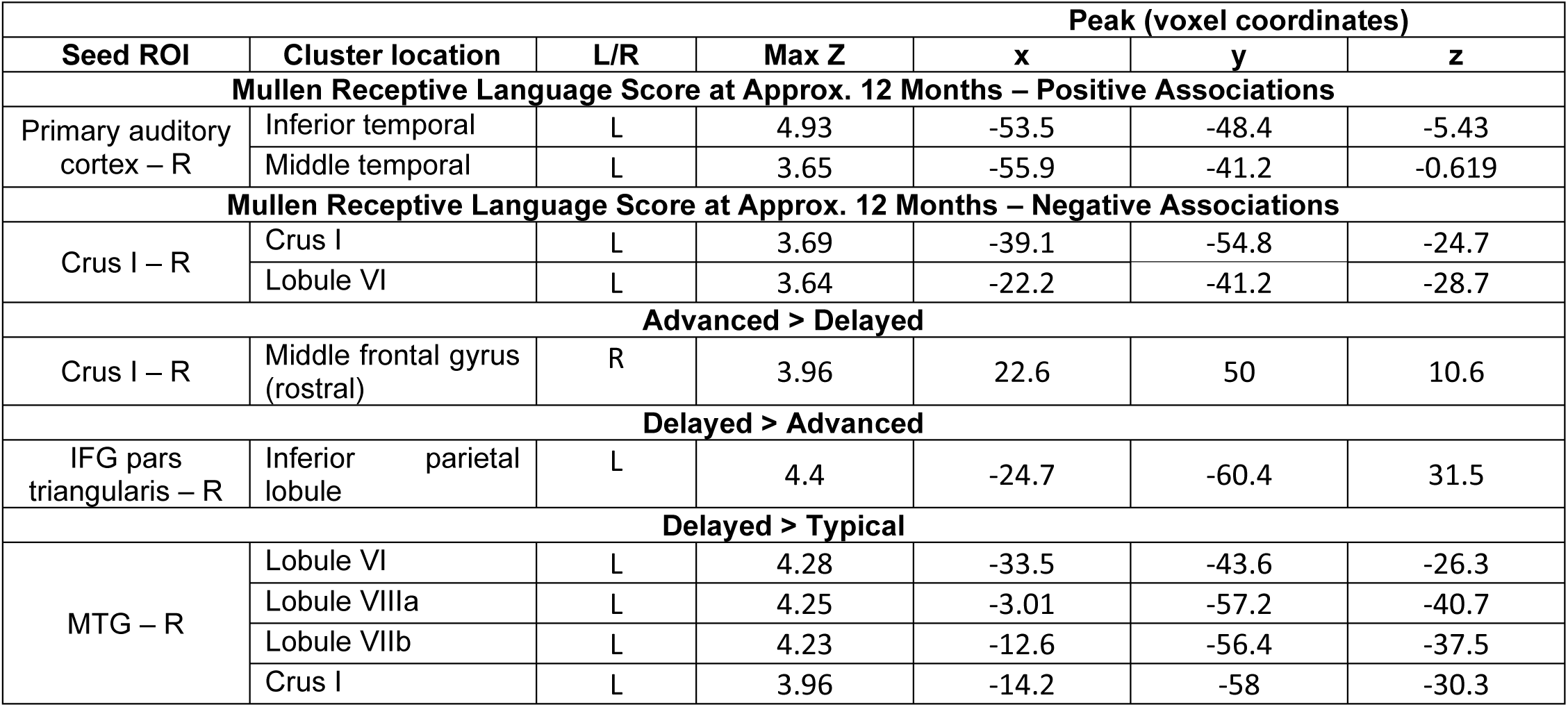
Coordinate table for peak clusters observed in language regressions and group differences. Bottom-up regressions for Mullen Receptive Language scores were masked by language network ROIs. Group contrasts were not masked. ROI = region of interest; L = left; R = right; IFG = inferior frontal gyrus; MTG = middle temporal gyrus.

The Delayed language group showed functional hypoconnectivity, relative to the Advanced group, between the right crus I of the cerebellum and a right prefrontal cluster localized to the middle frontal gyrus (Figure 5A). Connectivity of this cluster, which was stronger in the Advanced group, showed consistent positive correlations across the whole group with the MSEL expressive language subscale (*r* = 0.36, *p* = 0.002), MCDI productive vocabulary size (*r* = 0.33, *p* = 0.017), and the VABS communication subscale (*r* = 0.25, *p* = 0.049), all at approximately 2 years of age. This indicates that greater connectivity between right crus I and ipsilateral prefrontal cortex is predictive of better language skills over one year later.

**Figure 5.**
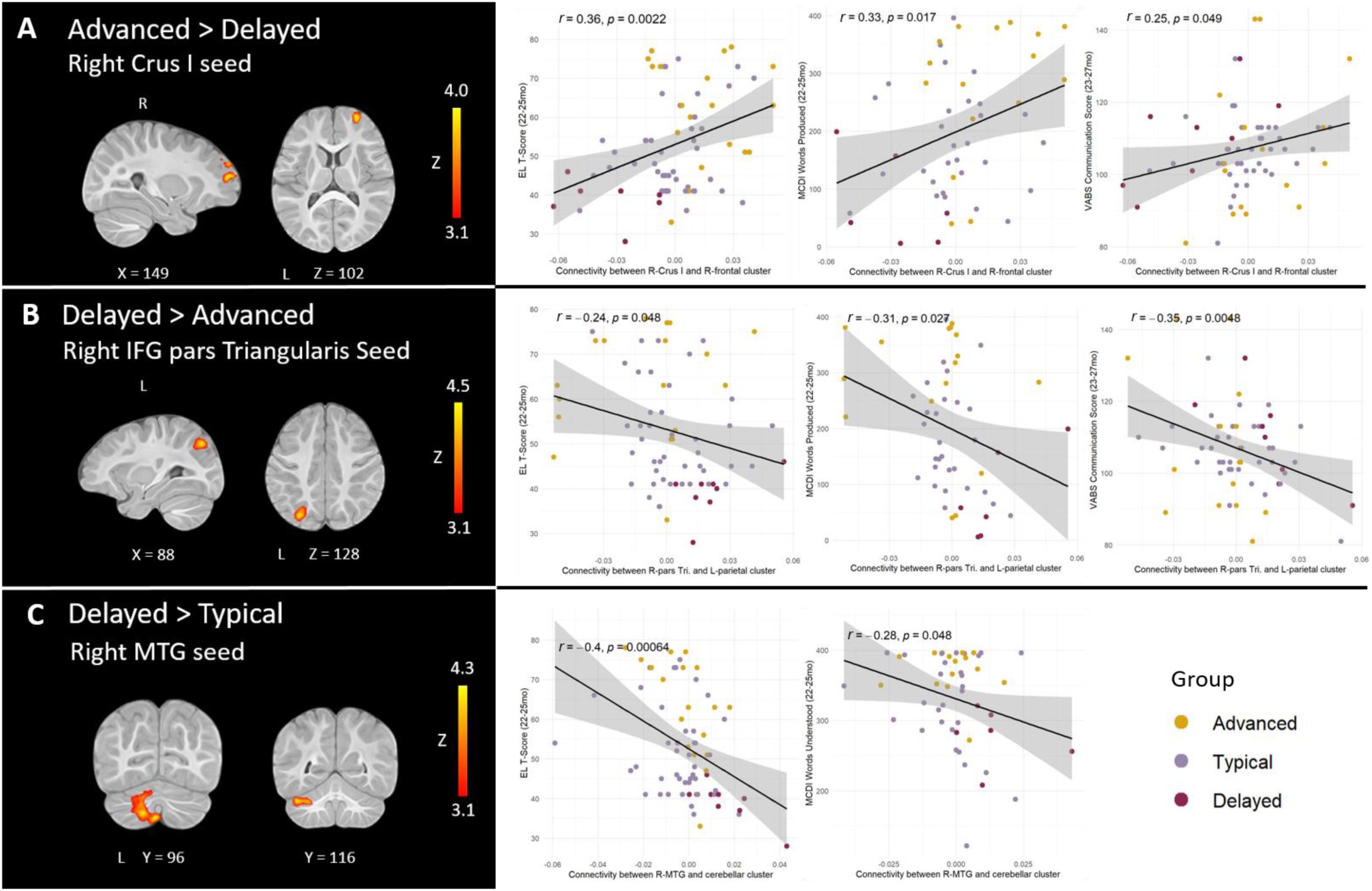
Differences in early connectivity between developmental language cohorts (left panel), and correlations with later language skills (right panel). Brain-behavior correlations were computed across all three language cohorts, regardless of the group comparison in which the difference was detected. All analyses regressed out the effects of site and age at the time of MRI scan. IFG = inferior frontal gyrus; MTG = middle temporal gyrus; EL = Mullen expressive language subscale; MCDI = MacArthur Bates Communicative Development Inventories; VABS = Vineland Adaptive Behavior Scales.

The Delayed group also showed hyperconnectivity relative to the other two groups. Language-delayed infants were characterized by atypically strong connectivity between the right pars triangularis of the IFG and the contralateral inferior parietal lobule relative to the Advanced group (Figure 5B), as well as atypically strong connectivity between the right MTG and a large left cerebellar cluster (spanning lobules VI, VIIIa, VIIb, and crus I) relative to the Typical group (Figure 5C). Since ethnicity was a potential confound for the Typical group, we confirmed that group differences in the Delayed vs. Typical contrast held by comparing parameter estimates from this cluster between Delayed vs. Typical-Hispanic (*t* = 3.97, *p* = 0.0007) as well as Delayed vs. Typical-Non-Hispanic (*t* = 5.21, *p* = 5.21e-5). Importantly, these clusters also showed remarkable associations with later language scores across several distinct language and communication assessments. The right pars triangularis–parietal connectivity cluster had reliable negative correlations with 2-year language scores, including the MSEL expressive language subscale (*r* = −0.24, *p* = 0.048), MCDI productive vocabulary size (*r* = −0.31, *p* = 0.027), and VABS communication score (*r* = −0.35, *p* = 0.0048). The right MTG–cerebellum cluster also had significant negative associations for connectivity strength, with MSEL expressive language scores (*r* = −0.40, *p* = 0.0006), and MCDI receptive vocabulary size (*r* = −0.28, *p* = 0.048). Therefore, stronger connectivity between these regions is predictive of worse language scores later in development.

## 4. Discussion

In this study, we examined neurodevelopmental profiles of language network functional connectivity throughout the first year of life with the goal of identifying normative developmental trajectories as well as early atypicalities that may predict later language delay. Using both whole-brain and pairwise analytical approaches, we found several consistent patterns of increasing intra-network synchrony and specialization throughout the first year of life, including age-dependent increases in both short- and long-range functional connectivity strength within the language network. Overall, we found strengthening of long-range connectivity between the IFG and primary and secondary auditory cortices, as well as with the thalamus. There were also significant increases in short-range connectivity between middle and superior temporal cortices, as well as within the cerebellum. By contrast, cerebellar ROIs consistently showed age-dependent decreases in connectivity strength with the MTG, and the thalamus also displayed decreasing functional synchrony with temporal ROIs as a function of age. Interestingly, stronger connectivity between right primary auditory cortex and a left temporal cluster significantly correlated with better receptive language scores at approximately one year of age, whereas greater connectivity between right crus I and left cerebellar language regions predicted worse scores. We also employed a behavior-driven clustering approach to identify three cohorts who displayed Advanced, Typical, and Delayed trajectories of receptive language development. Not only did these three cohorts reliably differ on an array of language measures, but they also displayed differences in language network connectivity that may be indicative of delayed language. These included underconnectivity between right crus I and prefrontal cortex, hyperconnectivity between right pars triangularis and left parietal cortex, and hyperconnectivity between right MTG and left cerebellum. Importantly, connectivity strength between these regions was reliably predictive of language development at two years of age across several independent language measures.

Here, we analyzed age-dependent changes in language network connectivity using two complementary approaches: a pairwise, targeted ROI-to-ROI approach, as well as a bottom-up seed-based connectivity approach. Across both approaches, we observed increasing specialization and synchrony between various modules of the language network. We replicated and extended some key maturational findings from prior work performed in a smaller longitudinal sample of 20 infants who had data at 1.5 at and 9 months of age (Liu et al., 2020). Consistent with Liu et al. (2020), our considerably larger sample also showed age-related FC increases between lower-level auditory areas (primary auditory cortex and thalamus) and the IFG, and age-related decreases between primary auditory cortex and ipsilateral pSTG. The bottom-up whole-network analysis indicated that both of these relationships may be especially strong for right primary auditory cortex. We also replicated decreasing thalamic connectivity with the pSTG with increasing age, a pattern that was not observed in infants with a familial history for ASD (Liu et al., 2020). Interestingly, a notable functional connectivity analysis of the language network in nearly 1,000 adults (Tomasi & Volkow, 2012) has demonstrated that, in addition to strong synchrony with its contralateral homolog and bilateral Broca’s area, Wernicke’s area also shows strong anticorrelations with primary sensorimotor areas, including primary auditory cortex. Taken together with our age-related results, we may conclude that temporal segregation between primary and higher-level auditory cortices may be a necessary step along the way to the adult functional network configuration.

We also observed some additional maturational changes that were not found in Liu et al. (2020), including broad, significant strengthening between the pSTG and IFG subregions. This encompassed patterns of increasing intrahemispheric and contralateral synchrony between Broca’s and Wernicke’s areas across the first year of life, which may lead to the bilateral synchrony previously observed across the core language network in adults (Tomasi & Volkow, 2012). Intrahemispheric connectivity between the pSTG and IFG is not observed at birth (Perani et al., 2011; Smyser et al., 2010), but is documented within the first year of life (Emerson et al., 2016). Although prior work has supported the notion of concurrent increases in long-range connectivity and decreases in short-range connectivity in typical development (Ciarrusta et al., 2020; Liu et al., 2020), our results suggest that short-range connectivity also increases, to some extent, between some key nodes of the language network whereby the pSTG showed significant inter- and intra-hemispheric strengthening with the MTG. This observation is difficult to contextualize, as prior work (Emerson et al., 2016; Liu et al., 2020; Tomasi & Volkow, 2012) did not examine MTG connectivity. However, given that information from both the middle and superior temporal gyri travels to dorsal premotor cortex via the superior longitudinal fasciculus (Friederici & Gierhan, 2013), some functional coupling between these regions would be expected. There was also a general trend of increasing intra- and interhemispheric connectivity within the cerebellum, supporting the cerebellum as an independent functional module at this age. This represents a novel contribution to the developmental study of the language network, as many prior investigations of this network have not considered the cerebellum despite its increasingly recognized role in language processing (LeBel & D’Mello, 2023), with the right hemisphere of the cerebellum being particularly implicated in language functions (Nettekoven et al., 2024).

Extending previously observed age-dependent decreases in thalamic-pSTG connectivity (Liu et al., 2020), our analyses also revealed that the thalamus shows broader functional decoupling from all temporal ROIs with advancing age, including decreases in thalamus-MTG and thalamus-primary auditory cortex connectivity. Additionally, we found strong evidence for functional connectivity decreases between all cerebellar ROIs and the MTG. Together, these relationships indicate that temporal language areas may undergo a general maturational trajectory of functional segregation from subcortical and cerebellar areas throughout infancy. Indeed, Tomasi & Volkow (2012) show that the cerebellum, thalamus, and STG are each found in distinct functional modules of the adult language network.

Notably, we did not observe consistent connectivity changes between homotopic Broca’s and Wernicke’s areas (i.e., changes in symmetrical interhemispheric connectivity) across the first year of life. Whereas prior work indicated that such symmetrical connectivity strength may peak for the IFG and STG around 12 months of age (Emerson et al., 2016), our findings are consistent with Liu et al. (2020), who found no significant changes in symmetrical connectivity for the IFG or pSTG between 1.5 and 9 months of age. Our results are also consistent with a more recent study conducted in the perinatal period that indicated interhemispheric functional symmetry may undergo significant strengthening through the first postnatal month only (Scheinost et al., 2022). Together, these findings indicate that the symmetrical connectivity observed shortly after birth for Broca’s (Perani et al., 2011) and Wernicke’s (Perani et al., 2011; Smyser et al., 2010) areas, as well as primary auditory cortex, which displays adult-like symmetrical connectivity at birth (Gao et al., 2015), may be relatively static after the perinatal period as these regions begin to undergo intrahemispheric strengthening.

We also examined how functional connectivity within the first year of life related to later receptive and expressive language development, as assessed at 12 – 15 months of age (as indexed by the MSEL). Although we detected no relationships with expressive language, stronger connectivity between right primary auditory and left temporal cortices during infancy was predictive of better receptive language in the second year of life. Although this relationship was detected across the first year of life, these two regions did not display any significant age-dependent changes in connectivity, suggesting that very early interhemispheric connectivity in temporal cortices might be predictive of these later language outcomes. Interestingly, a prior study of infants at high likelihood for ASD found that reduced network efficiency in temporal language regions (primary auditory cortex, STG, and MTG) was predictive of later autism diagnosis (Lewis et al., 2017). We also observed that greater functional synchrony between the right crus I and contralateral cerebellar language ROIs predicted worse receptive language. Curiously, connectivity between these regions appeared to strengthen across the first year of life, according to our age-related analyses. It is possible that, even if increases in interhemispheric connectivity between cerebellar language regions are normative, atypical overconnectivity may hinder the right crus I’s emerging role as a language related region, as already observed in infancy (Okada et al., 2021; Wagner et al., 2025).

Our behavioral clustering analysis identified three cohorts showing distinct receptive language trajectories in the present sample. These groups followed Delayed, Typical, and Advanced profiles of language acquisition within the first three years of life. Importantly, the proportion of infants designated as “Delayed” – about 12% – aligns well with a prior report that put the prevalence of language delay at approximately 1 in 10 children in the general population (Norbury et al., 2016), thus supporting the face validity of this clustering approach. Similar to prior work that employed this same approach in infants at varying likelihood for ASD (Okada et al., 2021), the three cohorts in the present study robustly differed on an array of language and communication measures that evaluated expressive language ability, vocabulary size, and more general communication skills. However, unlike Okada et al. (2021)’s findings, here we observed no group differences on a different measure of joint attention. We also found robust group differences on additional non-language measures indexing motor development and visual processing. This suggests that, in addition to identifying language-delayed participants, retrospectively partitioning participants by receptive language trajectory could also be useful for identifying infants who may experience broader developmental delays.

When we compared the Delayed, Typical, and Advanced language cohorts on language network connectivity, we found that the Delayed group was characterized by atypical hypoconnectivity between right crus I and right prefrontal cortex. Remarkably, this closely mirrored a finding previously observed in a sample of infants at varying familial likelihood for ASD (Okada et al., 2021) that was also clustered by receptive language trajectories. Older autistic children with language impairment also show underconnectivity between the right cerebellum and prefrontal cortex (Verly et al., 2014), suggesting that this atypical pattern may be persistent. Critically, in addition to extending prior work showing that right crus I–prefrontal connectivity may characterize populations with delayed or impaired language, we also found that, in our sample, the magnitude of this hypoconnectivity robustly predicted worse language skills later in development, at two years of age. These converging results, coupled with behavioral validation, suggest that right crus I–prefrontal connectivity may be an important early marker for language delay and a potential target for future investigations of early language network development. Indeed, the right crus I is commonly identified as a cerebellar region involved in language tasks (Nettekoven et al., 2024; Wagner et al., 2025). Prior work has also found right crus I hypoactivation during speech processing in HL infants (Wagner et al., 2025), in whom language delays are much more common than in the general population (Garrido et al., 2017). Clinical research has found that early damage to right crus I can even cause symptoms of dysfluency and reorganization of left frontal language regions in children (Riva et al., 2019). Whereas cortical language functions are left-lateralized, cerebellar language function is usually right-lateralized in adults and older children, reflecting the structural contralateral decussation of cerebro-cerebellar white matter tracts. Therefore, the favorable ipsilateral connectivity observed here is intriguing given that coactivation between the right cerebellum and left cerebrum is standard during language processing (Jansen et al., 2004). Although contralateral coactivation is normative in the adult network, the present results indicate that right-hemisphere prefrontal areas may also play a role in the cerebellum’s functional specialization for language earlier in infancy.

The Delayed group was also characterized by hyperconnectivity in two language regions. Importantly, these two hyperconnectivity patterns also presented significant associations with later language measures at two years of age, with stronger signatures of hyperconnectivity reliably predicting worse outcomes across all three groups. Connectivity between the right pars triangularis and left inferior parietal lobule was significantly stronger in the Delayed cohort. The inferior parietal lobule is divided into the angular and supramaginal gyri, both of which are thought to be involved in higher-order language functions such as semantics, phonology, and the integration of linguistic information between Broca’s and Wernicke’s areas via the dorsal language pathway (Coslett & Schwartz, 2018; Friederici & Gierhan, 2013). Given that right-hemisphere IFG subregions are not considered part of the canonical language processing network in adults (Friederici & Gierhan, 2013), overconnectivity with higher-order language areas such as the left inferior parietal lobule could reflect delays in interhemispheric specialization (i.e., excessive involvement of the right-hemisphere Broca’s area homolog). Indeed, prior work in typically-developing infants supports the notion that increasing interhemispheric segregation supports normative language acquisition, particularly for frontal regions (Emerson et al., 2016). Significant hyperconnectivity between right MTG and a substantial cerebellar cluster was also detected in the Delayed cohort. This left cerebellar cluster included language regions (crus I and lobule VI) as well as lobule VIIIa, a sensorimotor area (Stoodley & Schmahmann, 2010). This heightened right temporal-to-left cerebellar connectivity runs counter to the normative configuration of left temporal, right cerebellar specialization for language. Moreover, our age analyses indicated that functional synchrony between MTG and cerebellar language areas should significantly decrease throughout infancy. Therefore, persistent hyperconnectivity between these regions could be interpreted as a failure for functional specialization to develop in the Delayed group. Altogether, we identify three potential signatures of connectivity that could hold promise as early neural markers of language delay.

### 4.1 Limitations and Future Directions

Despite our large – for this population – longitudinal sample size, only a small subgroup of infants was identified as language delayed. Although the proportion of delayed infants within our sample was in line with prior behavioral literature on the prevalence of language delay in the general population (Norbury et al., 2016), this limited group size precluded further, more detailed analyses. Future studies examining functional connectivity associated with language acquisition should therefore aim to recruit a larger number of participants with suboptimal language trajectories. Additionally, none of our infants had a confirmed clinical diagnosis of language delay and, as a result, we were unable to determine how well functional connectivity profiles associated with language delay may map onto clinical diagnoses of any developmental language disorder. Future work should seek to validate our observed biomarkers in a well-powered sample of infants who have later, confirmed clinical diagnoses of language-related disorders, such as specific language impairment or ASD. It should also be noted that, although the Baby Connectome Project aimed to recruit a racially and ethnically representative sample (Howell et al., 2019), the final dataset used in this analysis was overwhelmingly white and non-Hispanic. Future studies should aim to recruit and retain a more representative sample to ensure that results are more generalizable. Larger-scale infant neuroimaging projects, namely the ongoing HEALthy Brain Child Development (HBCD) Study (Volkow et al., 2024) hold great promise for furthering this line of inquiry, including how atypicalities in functional connectivity of the language network may map onto the development of white matter structural connectivity across thousands of participants. Finally, a number of predictive classification models have used early resting-state fMRI data to classify infants based on clinical diagnoses of ASD (Emerson et al., 2017), age (Pruett et al., 2015), and other behaviors (see Scheinost et al. (2023) for review). The neural signatures identified in the present analysis may hold promise for improving such predictive models of language delay and related conditions, especially if used in combination with genetic and behavioral data that will be made available through the HBCD Study.

### 4.2 Conclusions

This study replicates and extends prior work on the functional maturation of the brain’s language network in infancy. We also identify several neural patterns associated with delayed language trajectories. Importantly, connectivity strength between these regions throughout the first year of life was robustly predictive of language skills at 2 years of age, across several distinct language assessments. These differences should be further investigated as potential biomarkers for early language delay to inform strategies for targeted interventions.

