## Supplemental Materials for "Language network functional connectivity in infancy predicts developmental language trajectories"

**SUPPLEMENT**

|  | **REGRESSION:** | **Age** | **MSEL** |
| --- | --- | --- | --- |
| *N* | | 154 | 113 |
| Age at Scan | | 6.4 ± 3.7 | 6.9 ± 3.5 |
| Sex | |  |  |
|  | Female | 91 (59%) | 65 (58%) |
|  | Male | 63 (41%) | 48 (42%) |
| Ethnicity | |  |  |
|  | Hispanic | 13 (8%) | 10 (9%) |
|  | Not Hispanic | 141 (92%) | 103 (91%) |
| Race | |  |  |
|  | White | 119 (77%) | 88 (77%) |
|  | Black | 5 (3%) | 4 (4%) |
|  | Asian | 4 (3%) | 3 (3%) |
|  | More than one race | 26 (17%) | 18 (16%) |
| Site | |  |  |
|  | UMN | 111 (72%) | 88 (78%) |
|  | UNC | 43 (28%) | 25 (22%) |
| Mean Abs. Motion | | 0.33 ± 0.31 | 0.34 ± 0.32 |
| Mean Rel. Motion | | 0.15 ± 0.06 | 0.15 ± 0.06 |
| Max Rel. Motion | | 2.31 ± 2.85 | 2.19 ± 2.50 |
| # ICA Components Removed | | 49.5 ± 18.9 | 48.4 ± 18.4 |

**Table S1.** Demographic distributions and MRI quality metrics for groups in the bottom-up regressions of age and language measures (receptive and expressive subscales). Abs= Absolute; Rel= Relative; MSEL = Mullen Scales of Early Learning; UMN = University of Minnesota; UNC = University of North Carolina; ICA = Independent Component Analysis.

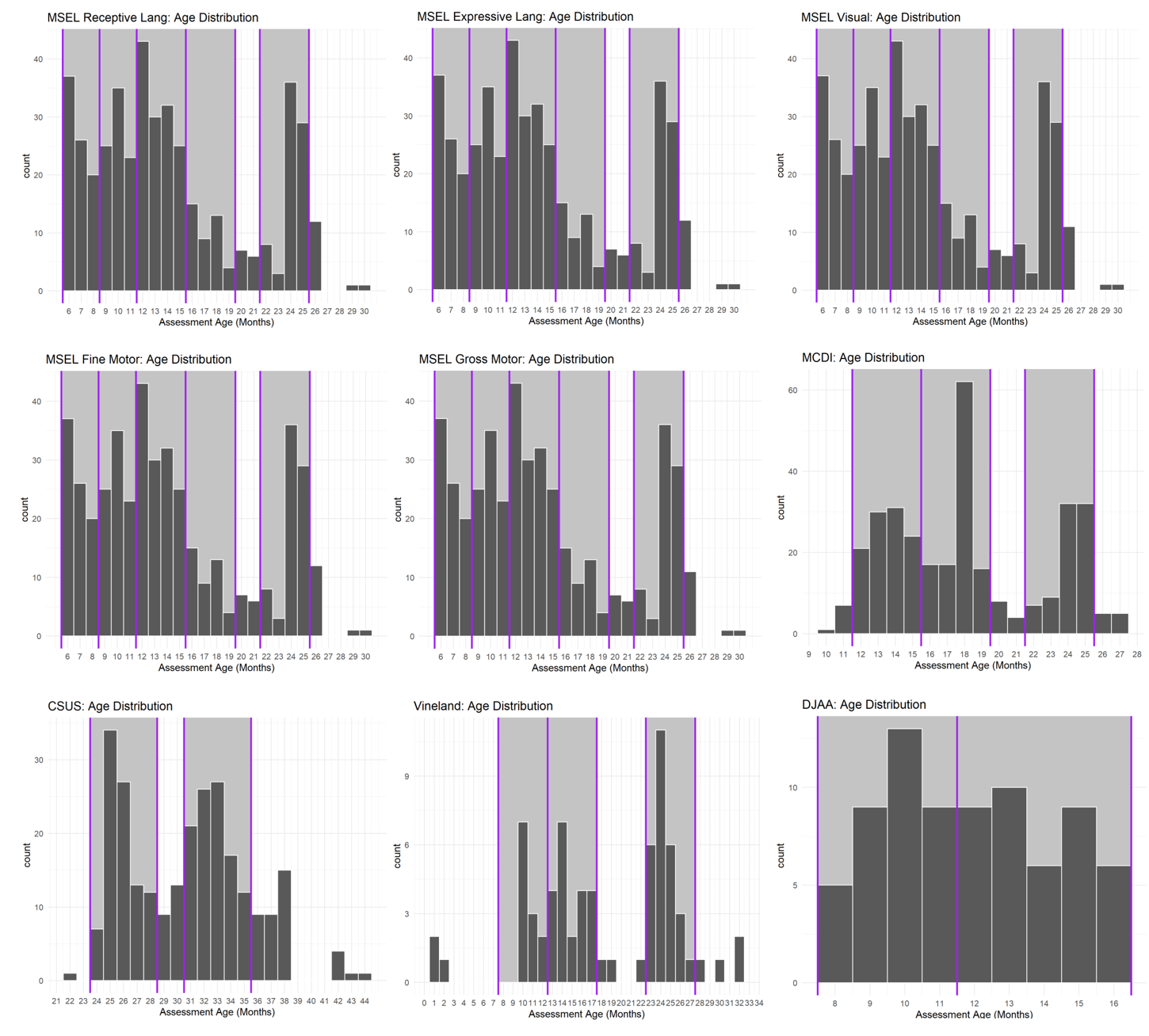

**Figure S1.** Age distributions of each behavioral measure. Age bins were spread evenly across the available ages as much as possible. Vertical purple lines indicate cutoffs for each age bin, and shaded magenta area indicates the distribution area included in each bin. From left to right, top to bottom: MSEL (receptive language, expressive language, visual reception, fine motor, gross motor subscales), MCDI (production and comprehension), CSUS, Vineland (all subscales), and DJAA.

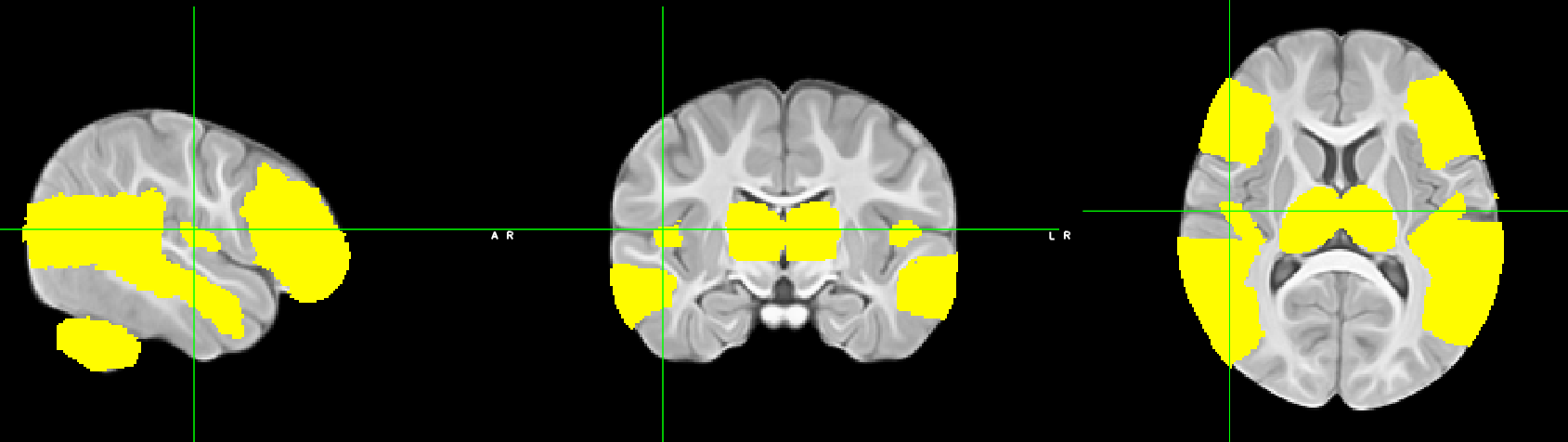

**Figure S2.** Mask of the language network. Because the Chen et al. (2022) atlas included only cortical gray matter parcellations, in order to include voxels within the underlying gray matter these cortical masks were derived from the Shi et al. (2011) 1-year infant atlas and subsequently warped to fit the Chen 12-month template.

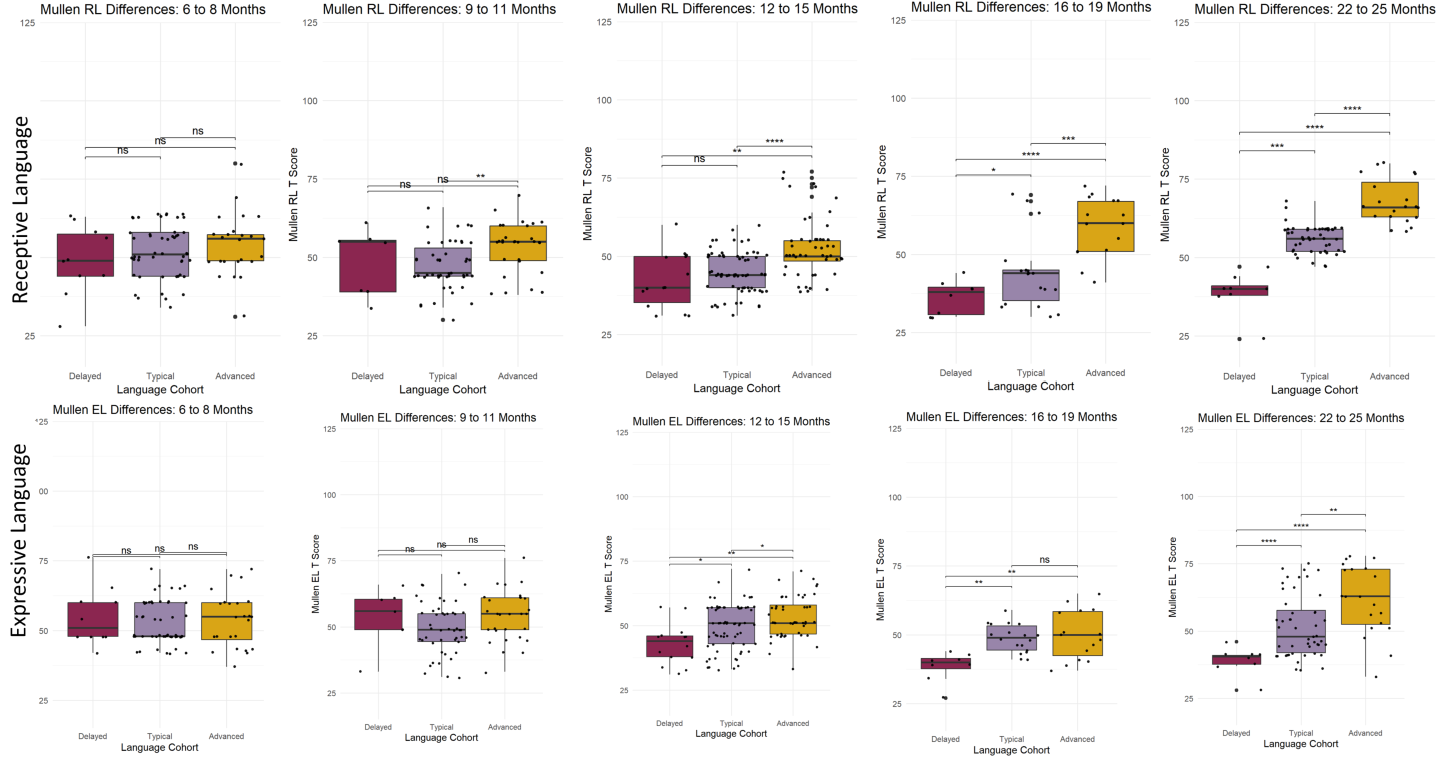

**Figure S3.** Cohort differences on the Mullen Scales of Early Learning (MSEL) receptive and expressive subscales across various age bins. RL= receptive language; EL = expressive language; ns = non-significant.
